# GENOAR: Global Engine for Navigating the Omnicell space via Agent-based Research

**DOI:** 10.64898/2026.10.15.759380

**Authors:** Hyojung Paik, Tae Lyun Ko, Dogyun Kim, Dongmin Shin, Marina Sirota, Boris Oskotsky, Tomiko Oskotsky, Taeyoung Lee, Hyeonjae Lee, Daewon Lee

## Abstract

Although generative models have revolutionized many fields, including genomics, building a scalable dataset compatible with artificial intelligence (AI) is essential yet underexplored. Here, we developed a Global Engine for navigating the Omnicell space via Agent-based Research (GENOAR) that covers approximately 54 million cells for diverse downstream tasks using AI agents and parallel computing.

GENOAR outperforms in three parts. Parallel computations accelerated the preprocessing of single-cell transcriptomic data; the other components enabled agent-assisted data storage and exploration for both AI models and humans. GENOAR involves two AI agents: an AI nomenclaturist ordered the unstructured labels of the single-cell transcriptomic profiles, and an AI librarian retrieved data. GENOAR presents transcriptomic profiles from more than 54 million cells from humans and mice in an analysis-ready format for humans and AI agents. AI agents in GENOAR exhibited successful retrieval performance compared with human experts. Using GENOAR, we elucidated progenitor cell-associated transcriptional signatures in native retinoblastoma and organoid environments. GENOAR was released and validated for the on-demand construction of single-cell atlases for metazoa. In conclusion, using GENOAR, the development of a single-cell transcriptomic atlas compatible with both AI systems and human researchers is feasible, scalable, and reproducible.

**Highlights:**

- Using AI agents and parallel computing, a single-cell atlas compatible with AI systems and human users can be feasibly developed and is inherently scalable as the dataset grows.
- The developed engine, GENOAR, enabled the establishment of single-cell atlases containing millions of cells with AI agents.
- On-demand construction of single-cell atlases is feasible and iteratively expandable.

## Introduction

Generative models, particularly large language models (LLMs), have markedly advanced many biomedical fields. For example, on the basis of data-rich pretraining using trillions of genetic variants, genomic LLMs are beginning to facilitate the functional annotation of previously uncharacterized variants[1]. Furthermore, advances in artificial intelligence (AI) agents have enabled the development of virtual labs that facilitate streamlined collaboration across interdisciplinary fields[2]. These transformative advances are fundamentally enabled by the availability of large-scale datasets organized by experts and shared through open science initiatives and public repositories, such as the Gene Expression Omnibus (GEO)[3]. Although millions of high-throughput molecular data points, including single-cell transcriptomics data, have been deposited in public repositories for transparency and reproducibility of data-driven studies, the scalable utilization of these data is still unfeasible. For example, in the first step of data reuse, public repositories are explored mainly through subjective inquiries, necessitating additional expert-driven data acquisition[4].

One promising approach to address this data readiness gap is large-scale, manual curation by human experts or consortium-level organizations. For example, the broad-scale application of single-cell RNA-seq (scRNA-seq) has enabled the collection of atlas-scale data such as the Human Cell Atlas (HCA), which encompasses millions of cells[5]. Effective foundation models[6, 7] in single-cell transcriptomics are built upon self-attention-based transformer structures and pretrained on reference datasets, including the HCA and CZ CELLxGENE collections (https://cellxgene.cziscience.com/), generated through consortium-like approaches. On the basis of the effectiveness of the pretrained LLMs in the field of single-cell transcriptomics, domain-adapted models for various fields, including cancer, have been developed[8, 9]. The scaling of both models and data is critical for constructing foundation models that can effectively elucidate latent biological interactions. However, structuring public scRNA-seq data to ensure its compatibility with AI models largely depends on human expertise, creating a scaling bottleneck as the data volume increases. Similarly, the scale of scRNA-seq atlases enable us to comprehensively understand diverse biological conditions including brain development[10] and diseases[11, 12], whereas models are constructed mainly on the basis of static data retrieved by human experts. Bulk-release paradigms in AI modeling and atlas-level explorations limit dynamic research, which could slow AI-driven discovery and the exploration of single-cell atlases. Although FAIR (findable, accessible, interoperable, and reusable) data principles[13] have promoted data availability, data reusability remains inconsistent, and the on-demand preparation of datasets for both AI systems and human experts remains immature.

Parallel computing technology, including compute unified device architecture (CUDA)-powered graphics processing units (GPUs), has enabled unprecedented advances in AI across diverse scientific disciplines[14, 15]. GPUs accelerate the training of large-scale AI models, including AI agents[16], through global memory, whereas the parallelization of multinode systems increases the flexibility for diverse tasks common in omics analyses. In our previous work, a dynamic workload strategy using message passing and workflow orchestration enabled the solution of otherwise unfeasible high-degree polynomial-time problems in biomedical fields within practical timeframes [17]. These computations are effectively scaled through distributed-memory parallelism. However, previous implementations in bioinformatics[17–20] have largely addressed isolated tasks, such as computational modeling, and the scalability of the data acquisition process remains unexplored. For example, modern generative language models such as InstructGPT[21] demonstrate that static corpora alone are insufficient for ensuring sustained performance, supporting the value of on-demand data acquisition at various time scales for model training.

scRNA-seq, which enables the characterization of different cell types and improves our understanding of disease pathogenesis, has elucidated tissue heterogeneity, lineage tracking, and disease mechanisms and may ultimately enable the development of personalized therapeutic strategies[22, 23]. Advancements in single-cell technology promote the development of diverse modalities and the integration of multiomics data[24, 25]. These breakthroughs in sequencing technology and LLMs have also raised new research questions, such as the feasibility of closed-loop data acquisition for AI models[26] and approaches for scaling training sets[27]. Methodologies capable of effectively harnessing, enhancing and adapting to large amounts of rapidly acquired scRNA-seq data for AI and human researchers must be developed. Although the automation of research with AI has been demonstrated, including idea generation, experiment execution, and manuscript preparation[28], data acquisition still relies mainly on query-based processes by human researchers, limiting the automation of data-driven biological research involving AI agents and humans. Although single-cell omics technology enables the characterization of key cellular properties across diverse biological contexts (e.g., diseases and tissues), the lack of automated pipelines for the analysis of large-scale scRNA-seq data and analysis-ready databases limit advances in single-cell biology.

Here, we developed a Global Engine for Navigating the Omnicell space via Agent-based Research (GENOAR, www.genoar.ai) for the on-demand development of single-cell transcriptomic atlases that are compatible with both AI and human interpretation. GENOAR retrieves scRNA-seq data considering the context of the corresponding biological experiment, including the age of the cells and specific disease. We were inspired by the concept-centered organization of biomedical language in the Unified Medical Language System (UMLS), which was developed by the National Library of Medicine, which integrates more than 2 million terms representing 900,000 biomedical concepts[29]. In our previous work, we used the UMLS to harmonize diverse biomedical terminologies on the basis of the underlying concept (i.e., meaning), thereby enabling large-scale biomedical analysis [30–33]. Building upon this concept-based framework, GENOAR systematically structures sample-and study-level metadata submitted to the GEO, such as tissue type, disease status, and experimental conditions, using both human experts and AI agents, including a retrieval-augmented generation (RAG) strategy and a model context protocol (MCP). Afterward, GENOAR retrieves public scRNA-seq data from the GEO, systematically annotates sample-level metadata, and validates its consistency with expert-curated metadata. In addition, using parallel computing technology, GENOAR presents preprocessed scRNA-seq results aligned with the structured metadata; these results are compatible with AI models via an application programming interface (API). The three key components of GENOAR (i.e., the AI agents, parallel computing technology and concept-based data annotation strategy) are released as a source code for the on-demand generation of single-cell transcriptomics atlases and were validated via the systemic on-demand generation of a mouse single-cell transcriptomics atlas. In addition, we validated the data of GENOAR by examining its transcriptional atlases. We identified transcriptional signatures in the tumor microenvironment[34]. Taken together, GENOAR provides atlases of more than 54 million cells from human and mouse samples and can seamlessly be integrated with AI agents in an inherently scalable manner.

## Results

### Transition from manual analysis to a scalable framework in single-cell transcriptomics

The construction of atlas-level databases of single-cell transcriptomic data (i.e., scRNA-seq) using public repositories has depended mainly on manual analyses by human experts. For example, Kim et al. successfully illustrated the dynamics of the expression of genes associated with disorder risk during development by integrating multiple scRNA-seq datasets[12]. The overall process of data integration by a human expert involves the piecemeal integration of data and the structuring of the metadata for each study on the basis of specific research interests, such as analyses of B cells across different tumor conditions (**Figure 1A**)[35]. However, *atlases* are broadly defined as *collections of maps*. In cartography, these typically depict various geographies and can provide multiple views of the same geography (e.g., topography, roads, political boundaries, demographics). Similarly, single-cell transcriptomics *atlases* ought to capture diverse properties across identical conditions; however, such integration is not scalable when performed manually.

**Figure 1.**
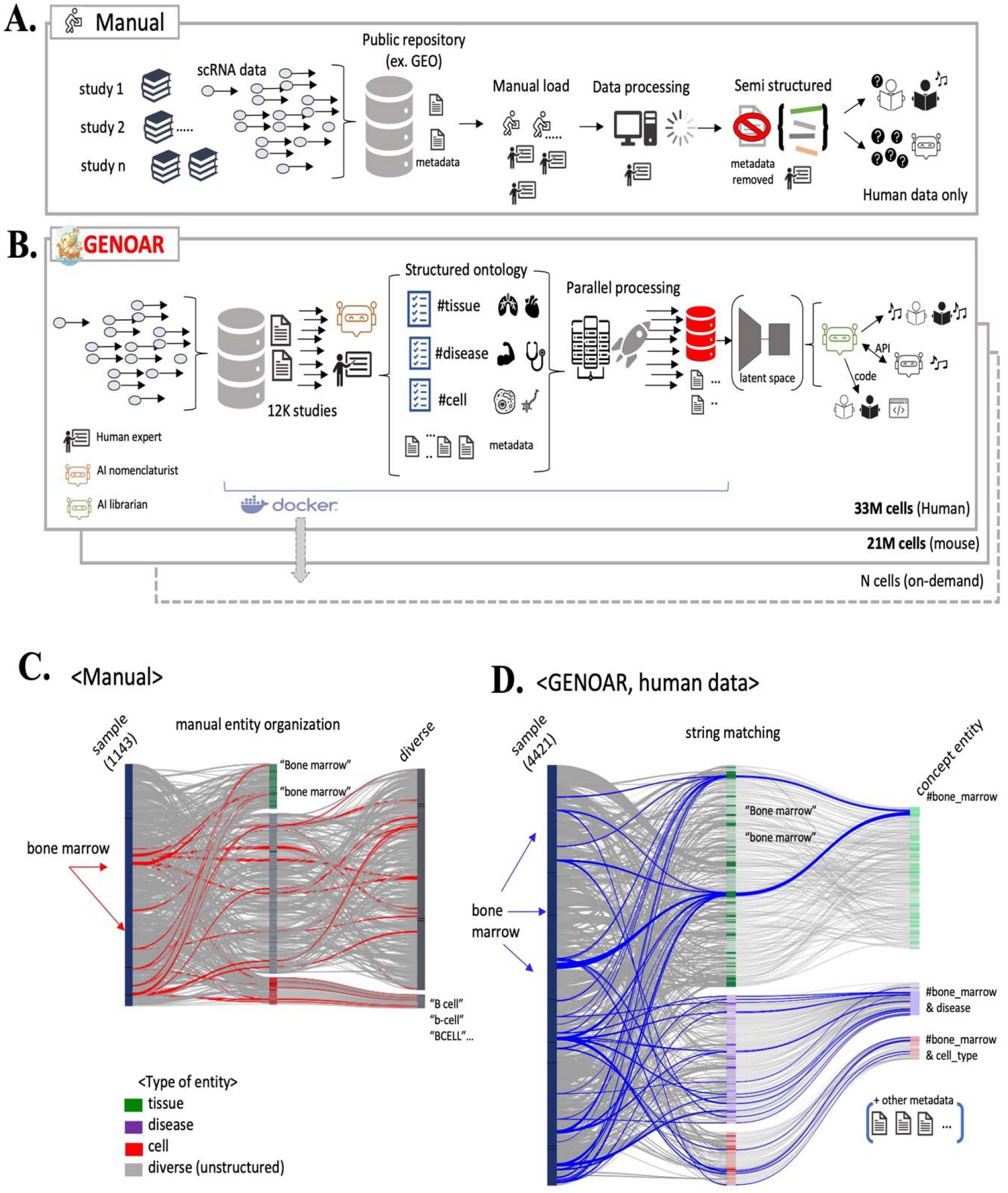
Construction of a large-scale single-cell transcriptomics atlas manually and with GENOAR. **(A)** Overview of a pipeline for the establishment of single-cell transcriptomic datasets manually by human experts. On the basis of a public repository of transcriptomic data, including the Gene Expression Omnibus (GEO), human experts curate data and perform preprocessing procedures. **(B)** In GENOAR, through the use of AI agents and parallel computing technology, human and mouse data from the GEO are systematically labeled at the concept level for each ontological entity and then preprocessed using parallel computing technology. **(C–D)** Flow of a human expert-guided organization of bone marrow samples (red lines) (C). GENOAR’s organization process for bone marrow samples (blue lines) was based on the AI nomenclaturist, which was guided by a human expert. In contrast to (C), the GENOAR system links additional metadata for each sample, including ages, sex and diverse conditions.

**Figure 1B** shows how GENOAR draws the analogous of atlas of single cell transcriptomics from the initial data labeling for diverse conditions, parallel processing of scRNA-seq data toward embedding of data label into latent space to present compatible with AI and human researchers. The pipeline of GENOAR starts with retrieving programmatic metadata from the GEO (https://www.ncbi.nlm.nih.gov/geo/) on the basis of predefined constraints, including the date (January 1, 2010 to August 31, 2025), species (*Homo sapiens*), and type of assay platform (10X Chromium, single-cell RNA-seq, and single-cell transcriptomics). The metadata for each study submitted to the GEO includes key sample features, including the tissue type, sex and biological state, which were voluntarily provided without a typical naming convention. On the basis of the concept (i.e., meaning) thesaurus used in the UMLS and AI disease nomenclature, GENOAR systemically structures the retrieved single-cell transcriptomic metadata at the sample level. As depicted in **Figure 1B**, the pipeline components were all containerized using Docker to ensure reproducibility, after which the same pipeline was used to construct single-cell transcriptomic analysis for mice (detailed in the Methods section). From this point, we utilize mouse data to validate the on-demand establishment of different datasets for analysis by GENOAR.

**Table 1** shows detailed statistics of the data in GENOAR obtained via the proposed pipeline (**Figure 1B**). From the initial retrieved metadata, more than 12,000 studies involving human data were identified. By filtering the sample data on the basis of number of reads (> 50 million) and file formats (e.g., fastq, fastq-load.py, and TenX), more than 4,000 samples comprising 33 million human cells were categorized in terms of the tissue type, cell type, and disease. As an ondemand build demonstration, the GENOAR pipeline was used to systematically categorize more than 2,000 samples consisting of 21 million mouse cells. We note that the data categories (e.g., tissue type, disease and cell type) in **Table 1** represent only a core part of the data entity. However, the diverse metadata of each sample were embedded in the latent space for the AI librarian agent using the RAG scheme. Among the three core data entities, tissue type (or disease) was designated as a required (non-nullable) field. Owing to the substantial tissue information among the metadata retrieved from the GEO, we determined that the tissue entity was a required field. Thus, most of the samples in GENOAR contain metadata for at least tissue information as well as other additional core data entities if available. For the disease name entity, the AI disease nomenclaturist (MCP-based agent) ameliorated classification of diseases using the corpus of UMLS concepts in **Table 1**.

**Table 1.** Statistics of the data in GENOAR.

| Overall fractions | *Homo sapiens* | *Mus musculus*<br>(on-demand build) |
| --- | --- | --- |
| Number of studies |  |  |
| Studies in public repository<sup>1</sup> | 12196 (by GSE<sup>2</sup>) | 8171 (by GSE) |
| Concept tagged<sup>3</sup> studies | 2914 (by GSE) | 3175 (by GSE) |
| Filter passed<sup>4</sup> studies (>100M, file formats) | 593 (by GSE) | 773 (by GSE) |
| Studies presented in GENOAR<sup>5</sup> | 385 (4,421 samples<sup>6</sup>) | 556 (2,854 samples) |
| Core data category in GENOAR<sup>6</sup> |  |  |
| Disease tagged studies (by GSE) | 111 (1,633 samples) | 28 (77 samples) |
| Tissue tagged studies (by GSE) | 385 (4,406 samples) | 556 (2,854 samples) |
| Cell tagged type studies (by GSE) | 36 (210 samples) | 60 (302 samples) |
| Total cells in GENOAR | 33,103,559 cells | 21,026,077 cells |

<sup>1</sup>In the present study, we used the Gene Expression Omnibus (GEO) dataset (https://www.ncbi.nlm.nih.gov/geo/, publication date in GEO from January 1, 2010 to August 31, 2025). <sup>2</sup>The identifiers of the GEO series (GSE); one GSE presents a research project involving a set of samples and relevant expression data. <sup>3</sup>Using the Unified Medical Language System (UMLS) of the National Library of Medicine (release version 2024 AB), unstructured descriptions of data submitted to GEO, including sample conditions, were converted into concept unique identifiers (CUIs). <sup>4</sup>From the CUI-tagged dataset of the GEO, systemic filters were applied for data quality (read count > 50 million; file format including fastq, fastq-load.py, and TenX; involving at least the CUI of tissue category or more). <sup>6</sup>Core data entities for annotating and retrieving samples in GENOAR.

To demonstrate the advantages of the systemic approach for harmonizing metadata via GENOAR, we retrieved an additional human dataset from the GEO through manual curation by human experts. As presented in **Supplemental Data 1**, human experts manually integrated the submitted single-cell transcriptomic metadata and preprocessed the data as presented in **Figure 1A**. Although 1380 samples covering more than 8.5 million cells were retrieved, the metadata of 235 samples did not include a standardized identifier of the Sequence Read Archive (SRR identifier of SRA, https://www.ncbi.nlm.nih.gov/sra) and were thus excluded from our systemic comparison. The categorization of the metadata of these 1143 samples (with SRR identifiers) in the human expert-derived integration (6.9 million cells) is shown in **Figure 1C**. We note that **Supplemental Data 1** was gathered for independent research purposes. Thus, the metadata submitted to the GEO have been reorganized and filtered mainly according to the research interests of the human experts, including cancer, resulting in a semistructured format of the data field, including post hoc augmentation of ‘Fetus label by manual (Yest/No)’, ‘Disease naming by manual (Cancer with abbreviations and manual typing)’. Therefore, as highlighted by the red lines in **Figure 1C**, single-cell transcriptomic datasets from the same tissue of origin, including ‘bone marrow’ and ‘Bone marrow’, do not have matching retrieval results. Moreover, most reorganized data entities remained unstructured (gray blocks in **Figure 1C**) and required a heuristic combination process. For example, the tissue information of one scRNA-seq dataset (SRR13724343) was manually labeled as a ‘cord’ sample, with the ‘Fetus label by manual’ field as ‘yes’, indicating cord blood containing fetal cells.

In contrast, **Figure 1D** depicts the programmatic organization of GENOAR on a larger scale (n=4421). As highlighted in blue, the diverse corpuses of bone marrow (BM, Bone Marrow, Bone marrow) are harmonized through one concept identifier, and the RAG-based AI librarian and MCP-based AI disease nomenclaturist provide additional metadata, including the disease and cell type, if available. In terms of mouse metadata organization, **Supplemental Figure 1** demonstrates that the on-demand, large-scale systemic organization of the submitted metadata of single-cell transcriptomic data is feasible using GENOAR. In addition, **Supplemental Figure 2** presents detailed comparison results between the manual naming of diseases and tissues in Supplemental Data 1 and the classification results of GENOAR using the UMLS corpus (details in Methods). All the manually assigned disease (and tissue) names of the collected single-cell transcriptomic data are successfully reorganized using the concept identifiers from the UMLS, demonstrating that GENOAR can programmatically organize disease- and tissue-related information of single-cell transcriptomic data.

### Contribution of AI disease nomenclaturist in GENOAR

The GENOAR pipeline can be used to analyze manually organized single-cell transcriptomic data from public repositories, including the GEO, and then validate these data through comparisons with classified tissue information of the samples. Metadata fields describing the tissue types were effectively distinguished using corpus-derived features and concept information from the UMLS, whereas the classification of disease-related metadata necessitated more sophisticated approaches because of the inherent hierarchical structure of disease ontologies. As we noted in previous studies, hierarchical disease classification is valuable for capturing disease similarity, which can facilitate the analysis of comorbidity patterns[32] and reveal shared biological mechanisms relevant to drug repositioning[36]. Thus, an explicit presentation of these disease relationships provides structured representations that seamlessly improve the performance of AI-based modeling by AI agents and human researchers.

We employed an MCP-based agent, named the AI disease nomenclaturist, to present these disease relationships and extract conceptual disease definitions from the disease-related fields via GENOAR. The role of the disease nomenclaturist in GENOAR is shown in **Figure 2A**. With the guidance of human experts, we designed an annotation process for disease classification on the basis of given disease fields, as described in the Methods section. In summary, a human expert designed the rule for the AI disease nomenclaturist and provided a reference for disease classification (https://uts-ws.nlm.nih.gov/rest), which is systematically accessible via an API. Diseases are organized in a hierarchical manner, in which categories at level 1 are subdivided into diseases at level 2. Among the 1,628 samples in the human dataset of GENOAR, our AI-based disease nomenclaturist identified 154 level-1 disease concepts, corresponding to 224 diseases at the level-2 subhierarchy. Similarly, the AI disease nomenclaturist identified 18 disease-related concepts from the mouse data of GENOAR via an on-demand process. We note that the disease level (i.e., level-1 and level-2) in GENOAR is determined by a human expert for convenience (details in the Methods).

**Figure 2.**
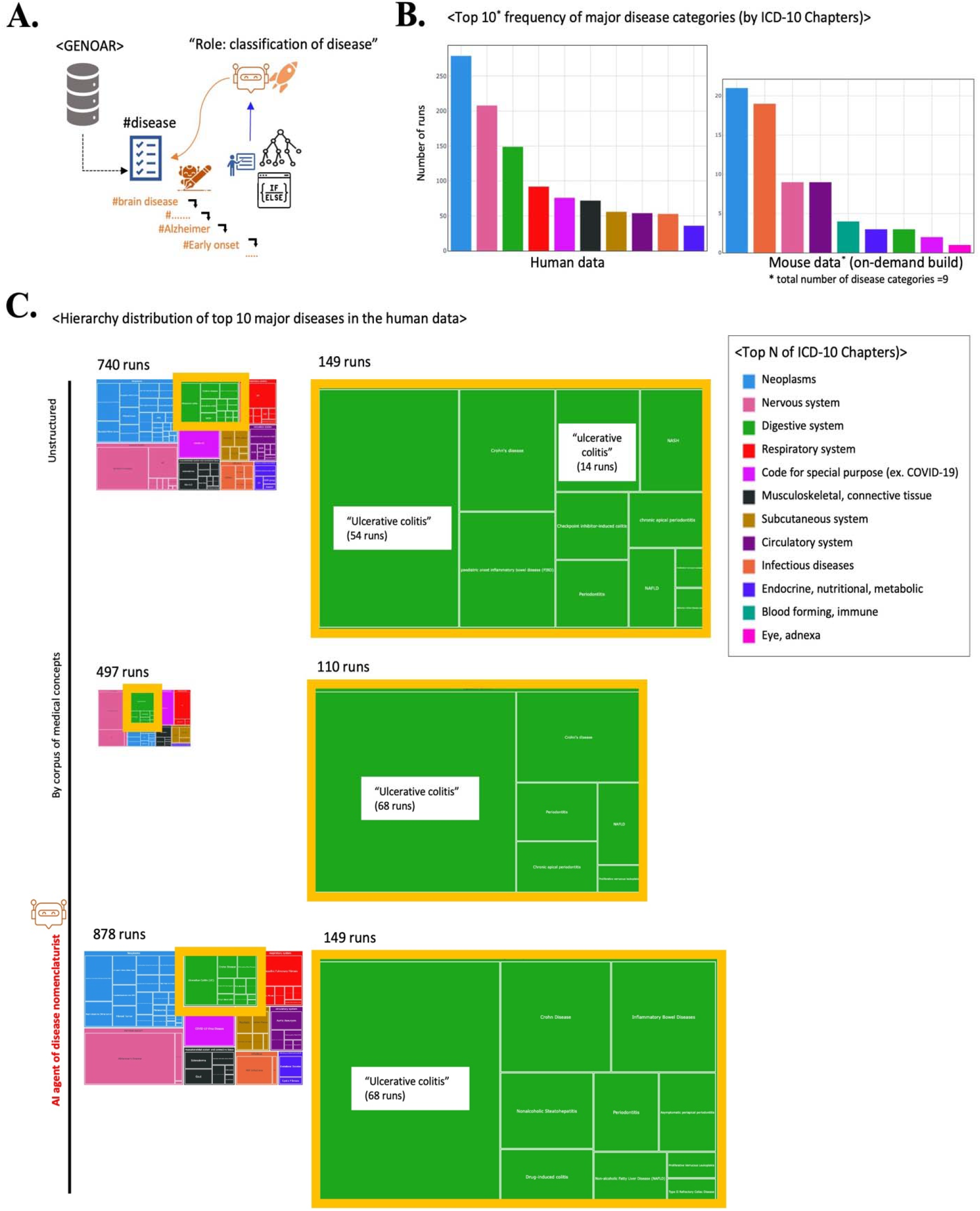
Systemic classification of a single-cell transcriptomics atlas using an AI disease nomenclaturist. **(A)** Role of the AI agent, named the disease nomenclaturist, in the systemic classification of the disease type. With the guidance of a human expert (see Methods for details), an MCP-based AI disease nomenclaturist classifies disease on the basis of the metadata submitted to the GEO. **(B)** Frequencies of the top 10 disease categories in humans and mice. **(C)** Tree-map visualization of the hierarchical distribution of the top 10 diseases in humans compared across three metadata annotation strategies: unstructured text (top), corpus-based medical concept matching (middle), and the AI disease nomenclaturist (bottom). Each block is colored according to the ICD-10 chapter (see legend). The yellow highlighted blocks indicate digestive system diseases, demonstrating that AI agent approaches consolidate heterogeneous disease representations into a single standardized concept, as exemplified by the unified identification of ulcerative colitis.

The frequencies of the top 10 disease categories in the human and mouse data from GENOAR are shown in **Figure 2B**. Disease categories were determined by mapping the disease concepts to the disease classifications from the World Health Organization (the International Classification of Disease-10^th^ release, ICD-10). The disease categories, such as neoplasms (shown in blue), include multiple specific disease names, including non-small cell lung carcinoma under the neoplasm chapter. Neoplasm (shown in blue) and Nervous system (shown in pink)-related diseases, including Parkinson’s disease, are frequently identified in the human samples retrieved by GENOAR, which effectively reflects overall research interests in the single-cell transcriptomics field. Interestingly, in the on-demand mouse data retrieval results, infectious diseases (shown in dark orange) and circulatory system diseases, including heart failure (shown in dark violet), frequently occur, indicating the scientific contribution of mouse-based approaches, likely because of the feasibility of tissue collection in mice. The overall disease distribution is consistent with current research interests in the single-cell transcriptomics field, ensuring that GENOAR retrieved single-cell transcriptomic profiles without bias.

On the basis of the confidence of the representativeness of GENOAR, we benchmarked the advantages of the AI disease nomenclaturist for the annotation of disease-related fields in the metadata through comparisons with other approaches, including manual naming and a UMLS concept-related corpus-based approach. The tree maps for the hierarchical distribution of the top 10 frequently-occurring disease categories in the human data by sample runs in **Figure 2B** are compared across three metadata annotation strategies: unstructured text usage (top), corpus-based medical concept matching (middle), and the AI disease nomenclaturist (bottom) (**Figure 2C**). The size of each block in the tree map indicates the number of samples (experimental run) for the corresponding disease category. The unstructured text-based approach directly uses the words provided in the disease-related field in the metadata, whereas the concept-matching strategy mainly uses the disease concept-related corpus (i.e., diverse synonyms of a given concept). As depicted in **Figure 2C** (top), the unstructured text-based approach yielded poorly organized sample runs for the same disease concept, including ulcerative colitis, from 740 disease field-matched samples in the metadata of GENOAR. Although the concept-matching strategy appropriately organized the sample metadata for the same disease concepts (middle of **Figure 2C**), the coverage of samples for corresponding disease-related concepts was insufficient, indicating substantial loss of disease-related information. Only 497 samples with disease-related concepts were hierarchically organized using the concept-matching strategy. However, when the AI disease nomenclaturist was used, 878 sample runs and corresponding diseases were systemically organized, and unified annotations were provided for each disease, including ulcerative colitis (bottom of **Figure 2C**).

**Supplemental Figure 3** presents further detailed results of the AI disease nomenclaturist using the human data and mouse data (on-demand build) of GENOAR. Similar to **Figure 2C**, **Supplemental Figure 4** shows a comparison of the hierarchical classification results of different diseases on the basis of mouse data using the above three strategies. As presented in **Supplemental Figure 4**, the AI disease nomenclaturist of GENOAR systematically classified diseases in a hierarchical manner with a minimum loss of sample information compared with that observed with the corpus-based medical concept-matching scheme. For example, for the hierarchical classification of blood- and immune system-related diseases (shown in turquoise), three systemic lupus erythematosus (i.e., lupus) samples were presented as a lupus-related mouse model, MRL/*pdcd*1^-/-^, which required manual interpretation by a human researcher. However, the AI disease nomenclaturist classified those three samples as lupus-related samples[37]. Although our disease nomenclaturist was guided by a human expert for the systemic classification of different diseases, **Figure 2** and **Supplemental Figures 3–4** present promising results for large-scale hierarchical disease classification.

### Evaluation of the ability of the AI librarian of GENOAR to navigate single-cell transcriptomic atlases

On the basis of tissue- and disease-related information from the analysis-ready single-cell profiles of humans and mice, we constructed a RAG-based AI agent (i.e., an AI librarian) to assist in the exploration of single-cell transcription profiles with GENOAR. The role of the AI librarian is to assist with metadata exploration within GENOAR. As depicted in **Figure 3A**, in addition to the structured ontology of the disease, tissue and cell type, diverse fields in the metadata of the single-cell transcriptomic profiles—including sex, age, treatment, perturbation, and behavior—are also embedded in the latent space of the language model used in the RAG framework of the AI librarian (details in the Methods section). We examined four language models, namely, PubMedBERT[38], MiniLLM (https://huggingface.co/MiniLLM), BioBERT[38], and SapBERT[39], and compared the retrieval performance of the AI librarians with that of human expert-based retrieval on the basis of synthetic queries using the human and mouse data (**Figure 3B**). Performance was evaluated on the basis of the retrieval-augmented generation assessment (RAGAS) score. The RAGAS score measures the context precision and context recall of the retrieval performance and the mean average precision (MAP) of the relevance of the generated answer. On the basis of the RAGAS score, we employed the top-ranked language model as the AI librarian in GENOAR. In addition, the normalized discounted cumulative gain at K (nDCG@K) was used to measure the effectiveness of the prioritized ranks of the retrievals by the AI librarian. In **Supplemental Data 2,** the details of the synthetic queries and expert-answered references are presented for the human and mouse data in GENOAR.

**Figure 3.**
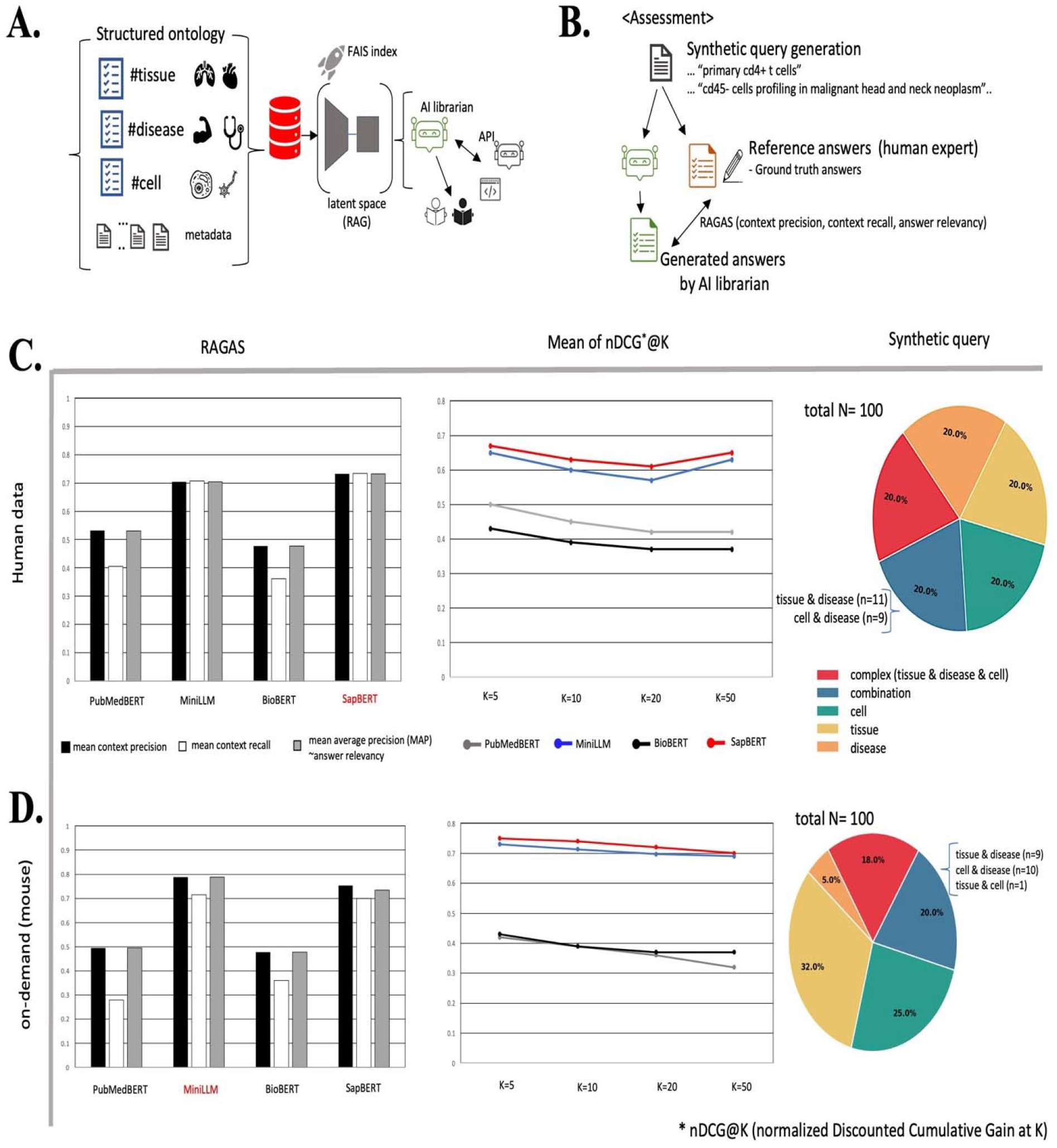
Evaluation of the retrieval performance of the AI librarian in GENOAR. **(A)** Role of the AI agent, named the AI librarian. With FAISS index (see Methods for details), an RAG-based AI librarian navigates the metadata of the samples of GENOAR. **(B)** Evaluation of the performance of an AI librarian using reference answer sets via retrieval-augmented generation assessment (RAGAS). **(C–D)** Evaluation of the performance of the AI librarian with human data (C) and on-demand mouse data (D). We established the RAG-based AI librarian using four models, namely, PubMedBERT, MiniLLM, BioBERT and SapBERT. We then employed the best-performing models in GENOAR on the basis of the results of the RAGAS. The pie charts present the proportions of categories of synthetic queries for each dataset (i.e., human and mouse data). Human expert-assigned reference answers are provided in **Supplemental Data 2**. These reference answers are used as the ground-truth set.

The retrieval performance and an overview of the synthetic queries (total of 100 queries) for the human data are presented in **Figure 3C**. In addition, we evaluated the performance of the AI librarian with the mouse dataset similarly (**Figure 3D**). First, we calculated the context precision and context recall for each query; then, the means of the context precision and context recall were determined. The context precision measures the proportion of retrieved documents (i.e., relevant answers of the AI librarian) that are actually relevant to the query on the basis of the reference answers of the human experts (**Figure 3B**). The context recall for each query is the proportion of reference answers that are successfully retrieved among all the documents retrieved by the AI librarian. The mean average precision (MAP) is a measure of retrieval relevancy and is calculated by averaging the precision scores at relevant positions across all queries. Here, averaging the precision scores involves calculating the precision at each rank position where the document, which is the relevant single-cell data (i.e., reference answer of the human expert), appears, after which the mean of these values is taken. The normalized discounted cumulative gain at K (nDCG@K) quantifies how well the relevant documents (i.e., reference answers of the human experts) were ranked among the top K results of the AI librarian in the search. Overall, when these measures are close to 1.0, the AI librarian is considered to appropriately guide the exploration of single-cell transcriptomic profiles in GENOAR. On the basis of the measures of the four language models, SapBERT was used to establish the AI librarian of GENOAR for 33 million human cells (mean context precision =0.732; recall = 0.734; MAP =0.732; mean nDCG@K=0.67 (K=5)) (**Figure 3C**). With respect to the exploration of 21 million mouse cells, the MimiLLM-based AI librarian outperformed the other models (mean context precision =0.787; recall = 0.714; MAP =0.787; mean nDCG@K=0.75 (K=5)) (**Figure 3D**). While the synthetic queries for human data included equal numbers of cell-, tissue-, and disease-related answers, we amended the proportion of disease-related queries among the synthetic queries for the mouse data while considering the sparseness of disease information for mice, as presented in **Table 1**. In conclusion, although the reference answers are inherently subjective because of the reliance on individual human expert judgments, overall, the retrieval performance of the AI librarian of GENOAR is similar to that of the human experts.

Thus, we analyzed the exploration of metadata on the basis of standardized disease- and tissue-related information and an AI librarian. While metadata standardization enhances data organization, its utility depends mainly on the availability of raw data for downstream use, considering the compatibility of such data with both AI agents and human researchers. On the basis of our previous work, we established a parallel computing pipeline to preprocess the single-cell transcriptomics data retrieved by GENOAR (**Table 1**). We employed a multinode computing system consisting of 1,488 CPUs and a dynamic workload scheme using Snakemake (https://snakemake.github.io/) to construct a reproducible and scalable single-cell atlas. In the current version of GENOAR, our parallel computing pipeline has processed and integrated more than 54 million cells from more than 1,200 studies, stored in an analysis-ready format (H5 format) and fully accessible via an API. Although GENOAR encompasses more than 2 petabytes of raw data, the Dockerized pipeline is compatible with diverse computing environments, ranging from local desktops to high-performance computing (HPC) clusters, allowing on-demand construction of datasets for all researchers (https://github.com/cxinsys/genoar).

On the basis of these promising results, GENOAR was established as a web resource (www.genoar.ai). **Figure 4A** shows the web interface, which mainly supports human researchers by showing the overall statistics of the data. The download section of GENOAR presents the core metadata, including the cell type-, tissue- and disease-related fields covering more than 54 million cells, which enables programmatic retrieval and is compatible with AI models via an API. Examples of the use of GENOAR via an AI librarian (**Figure 4B**) and the search results of a human researcher using a web interface (**Figure 4C**) are presented. Taken together, using AI agents and parallel computing technology, GENOAR presents preprocessed datasets covering more than 54 million cells, allowing the implementation of diverse downstream tasks and demonstrating compatibility with AI models.

**Figure 4.**
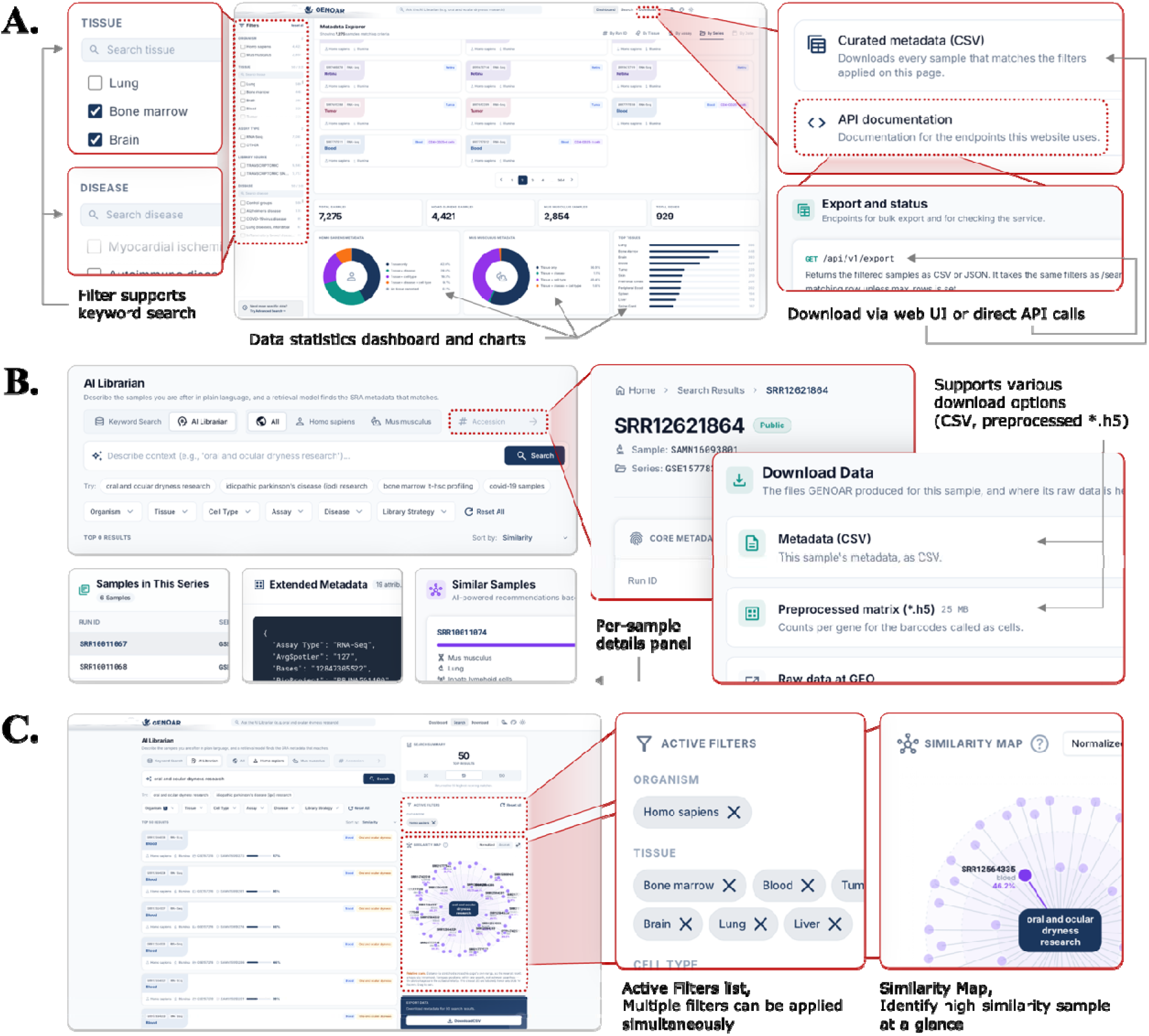
Web feature of GENOAR. **(A)** Dashboard of GENOAR for the first exploration. **(B)** Example of exploration results of the queries using an AI librarian and user-defined inquiry. **(C)** Functional features (i.e., additional filtering and similarity relationships) based on the inquiry results of GENOAR.

### GENOAR enables the construction of multicondition, multitissue single-cell atlases

Next, we validated the performance of the AI agents (disease nomenclaturist and librarian) and the reproducibility of the single-cell transcriptomic atlases via a systemic approach. To examine the validity of GENOAR, we conducted an integrative analysis using a dataset comprising multiple tissues and cancers. To address the described analog of single-cell transcriptomic *atlases*, we compared diverse tissue-related single-cell transcriptomic profiles with profiles representing diverse conditions across matched tissues. Using the human data in GENOAR, we identified 959,898 cells from tumors, control pairs (CPs) of tumors, and healthy controls from six tissues, namely, the retina, oral mucosa, myometrium, dura of the brain, colorectal tissue and bone cartilage. Although GENOAR contains considerable cancer data, we included only paired samples from solid tumors and matched CPs to construct single-cell *atlases*. To select healthy controls for each tissue, we selected samples from non-tumor-relevant studies using the standardized metadata in GENOAR (details in Methods). In total, thirteen studies (86 experimental runs of samples) were identified, after which we subsequently implemented the preprocessing procedure described in the Methods section, resulting in the generation of 950 thousand cells using a GPU-accelerated process (details in the Methods section). The performance of the GPU-based analysis pipeline was 732.4 times faster than that of the CPU-based process for these 950 thousand cells (**Supplemental Data 3, Supplemental Data 4)**.

To evaluate the preprocessing results of these 950,000 cells from the GPU-accelerated pipeline, we visualized the transcriptional features of the atlases cells using Unsupervised uniform manifold approximation and projection (UMAP) and t-distributed stochastic neighbor embedding (t-SNE). UMAP and t-SNE clustering analysis of the cells revealed results consistent with biological knowledge; namely, the transcriptional profiles of each cell are clustered by the corresponding tissue type (**Figure 5A**). The UMAP approach preserves the global structures of the clustered groups, and clusters from distinct tissues are clearly distant in the projection space (the UMAP plot in Figure 5A). Although fewer global relationships across tissues were preserved in the t-SNE results (left panel of Figure 5A), t-SNE-based clustering focuses on local structures. However, in t-SNE, cells originating from the same tissue type remain closely clustered, enabling the identification of diverse conditions associated with each tissue type. Thus, to address differences within each tissue across conditions, we mainly utilized the t-SNE clustering results in the subsequent analyses.

**Figure 5.**
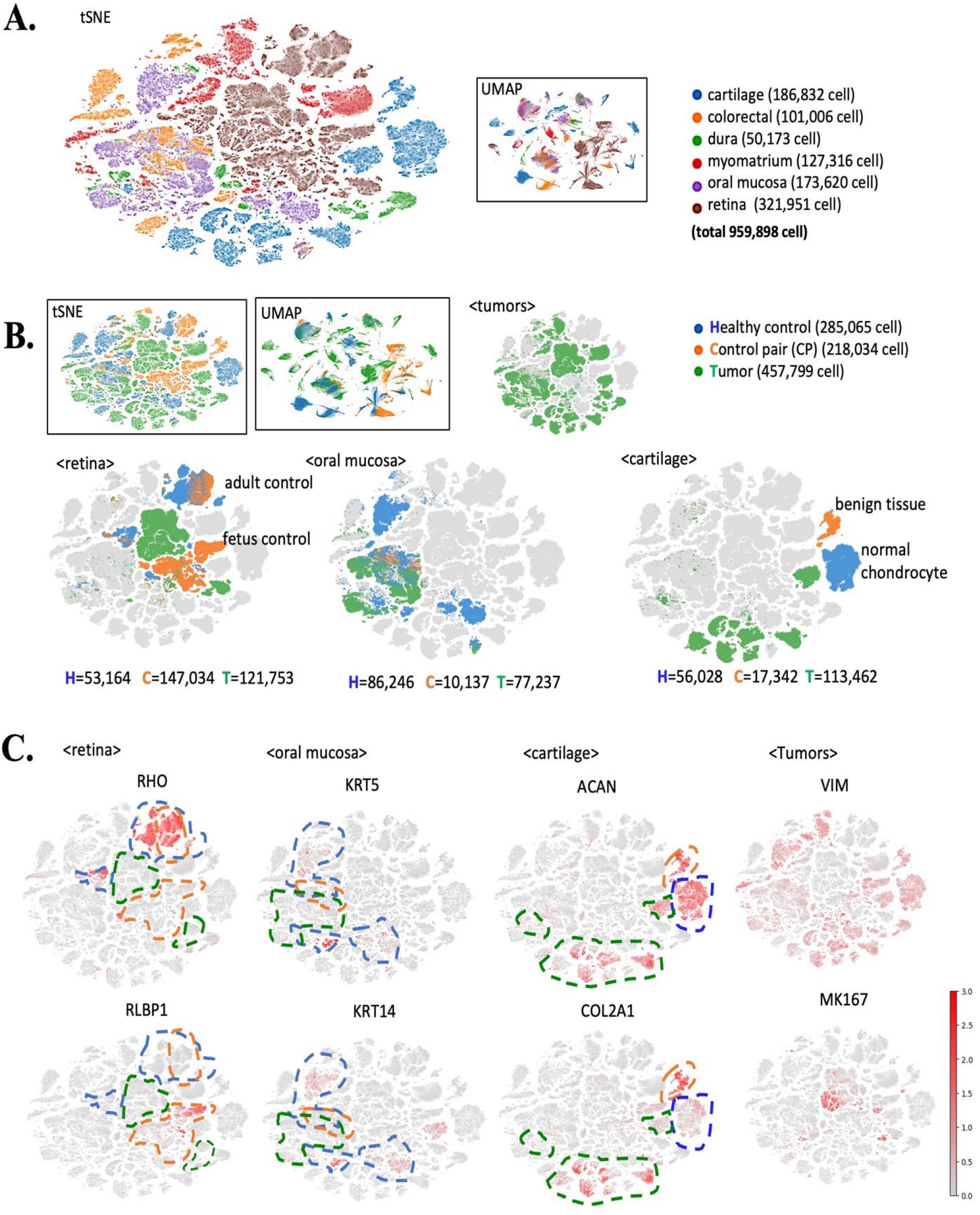
Identification of tumors, healthy controls and matched control pairs (CP) in GENOAR. **(A)** Results of dimensionality reduction and clustering of 959,899 cells derived from six tissue types (cartilage, colorectal, dura, myometrium, oral mucosa, and retina) using t-SNE and UMAP. **(B)** Featured projection of cells based on the identified tissue conditions, including healthy controls (blue), CPs (orange), and tumors (green). Detailed numbers of cells for each condition and tissue (retina, oral mucosa, and cartilage) are presented. **(C)** Intensity of the transcriptional signatures of canonical genes across tissue types and conditions. All cells and transcriptional signatures were processed through quality control and normalization steps (details in the Methods section) using the GPU-implemented pipeline.

**Figure 5B** presents the biological reliability across different tissues, including those from healthy controls (blue), CPs (orange), and tumor (green) tissue samples. For example, clustered cells from the retina were locally dispersed according to the tissue condition, including healthy controls (n=53,164), CP tissue (n=147,034), and tumors (n=121,753). The scRNA-seq data of the retina cells are based on three studies[40–42], including studies on retinoblastomas (RBs) and stem cell-induced tumor organoids. The CPs in these three studies involved diverse states, such as normal adult and fetal tissues. Among retina cells under CP conditions, cells from healthy adult donors were clustered with cells from deceased donors from other independent and noncancer studies (blue), whereas the remaining CP cells from distinct developmental stages (early trimesters) were clustered separately near healthy controls and tumor tissues. Thus, CP cells are clustered separately according to biological differences while preserving the unique features of the tissue of origin, allowing us to be confident in the integrative data provided by GENOAR. Similarly, the data from oral mucosa and cartilage tissues in the CP, healthy, and tumor conditions confirmed that GENOAR provides characteristic transcriptional profiles under diverse biological conditions within matched tissue atlases (oral mucosa = middle plot in Figure 5B; cartilage = the left plot in Figure 5B). The t-SNE results for the other three tissues, namely, the colorectal, myometrium and dura, confirmed that transcriptional profiles in different tissues across various conditions are presented properly in GENOAR (**Supplemental Figure 5A**). The detailed t-SNE clustering results in **Supplemental Figure 6** confirm the batch-effect correction of the analyzed data, including diverse CPs, tumors and healthy controls for each study.

Encouraged by the concordant clustering trends aligned by conditions and tissues, we next examined more detailed transcriptional signatures. The expression of canonical genes in the retina, oral mucosa, cartilage, and overall tumor are shown in red in **Figure 5C**. In addition, we marked borderlines in three colors to indicate the conditions of the corresponding tissues (blue = healthy control; orange = CP; green = tumor). For instance, rhodopsin (RHO) and retinaldehyde binding protein 1 (RLBP1) are essential genes in the retina [43] and are expressed exclusively in retinal cells (the left t-SNE plot in **Figure 5C)**. The exclusive expression of KRT5[44] and ACAN[45] in oral mucosa or cartilage allows us to be confident in the biological reliability of the GENOAR data. The data in **Supplemental Figure 5B** also support that the data of GENOAR recapitulate the known transcriptional signatures of the corresponding tissues. More specifically, the canonical genes of specific tissues are expressed mainly in healthy controls and CPs. In addition, tumor-related canonical genes, including VIM, MKI67, and VEGFA, broadly aligned with the cells derived from tumors, although the separation revealed a soft boundary, indicating the heterogeneity of the cancer microenvironment (**Figure 5C** and **Supplemental Figure 5C**). Although we explored a limited number of tissue-related genes, these results suggest that GENOAR successfully recapitulates the transcriptional signatures of these tissues across the various conditions at the single-cell level.

Taken together, the results indicate that GENOAR depicts diverse transcriptional profiles across matched conditions within the same tissue, analogous to geographic *atlases*.

### GENOAR prioritizes single-cell transcriptional signatures of RB

We next examined whether the *atlases* of GENOAR can be used to validate biological findings. We investigated integrated single-cell atlases associated with RB, a rare and life-threatening type of pediatric cancer. As many cells under RB conditions were available (Figure 5), we further analyzed these cells. As shown in **Figure 6A** and **B**, the retinal cells from healthy controls, CPs and tumors were obtained from three studies, namely, GSE147979, GSE174200 and GSE166173. We note that thousands of cells were filtered for this analysis by applying more strict preprocessing pipeline, as described in the Methods section. Thus, 234,689 cells were included in this analysis (321,951 cells in **Figure 6A**). Cells of CPs are involved in diverse conditions, such as fetal retina cells from GSE166173 and normal adult donor cells from GSE174200 (**Figure 5B**). In addition, the tumor cells shown in **Figure 6A–B** were primary RB [41] and stem cell-derived organoids of RB[42], which partially recapitulated the native tumor microenvironment (TME)[46]. However, RB cells from primary tumors and retinal tumor organoids were closely clustered, indicating broadly similar transcriptional profiles between native tumors and RB organoids (green in **Figure 6B**), warranting further analysis.

**Figure 6.**
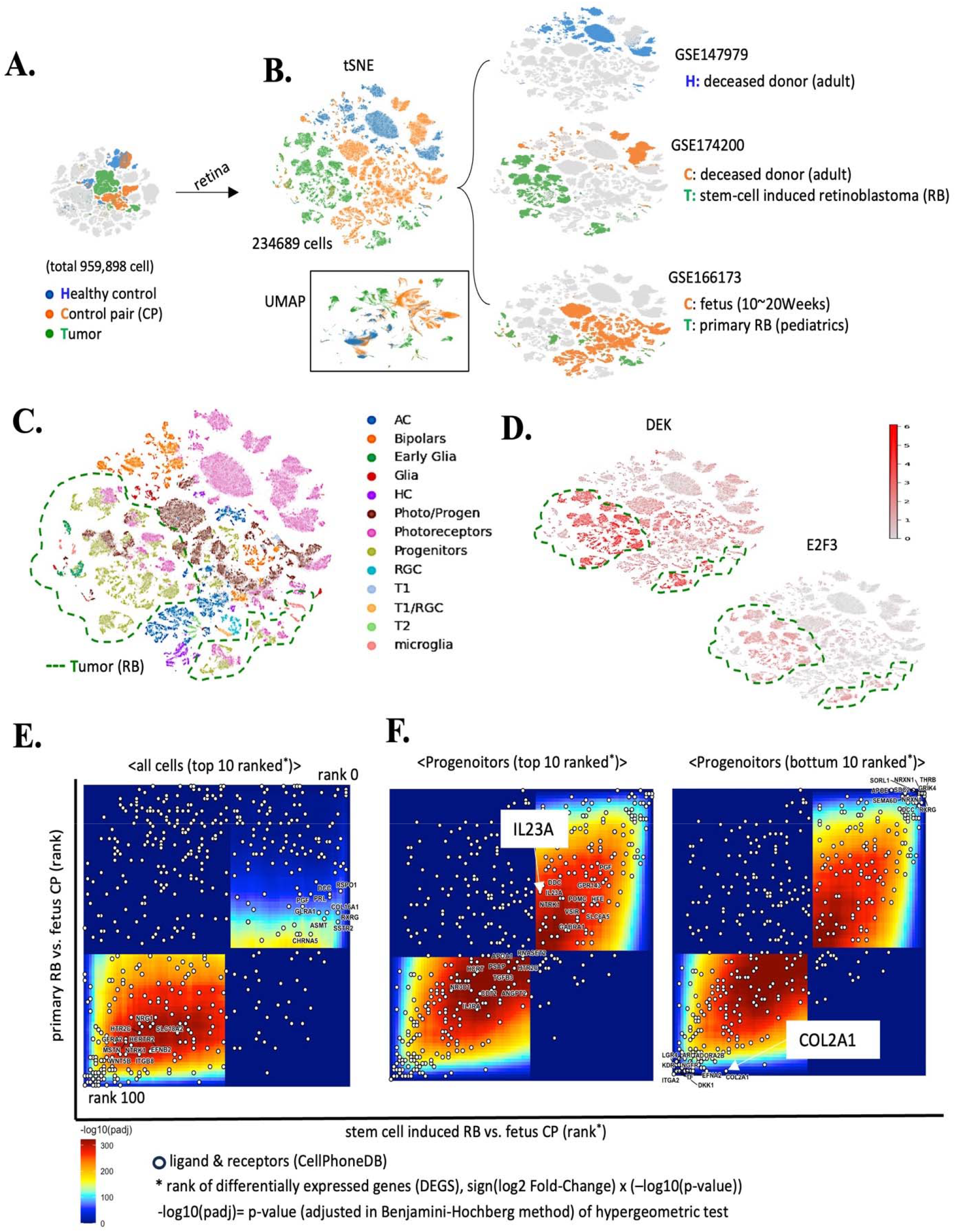
Identification of transcriptional signatures of retinoblastoma (RB) from primary tumors and organoids. **(A)** t-SNE plot of the retinal cells by condition, including cells from healthy controls (blue), CPs (orange) and tumors (green). (**B)** Results of re-clustering of cells from the retina. **(C)** Cell types were identified using the CellTypist method. **(D)** Expression of known retinoblastoma (RB) genes, including DEK and E2F3. The green lines indicate cells from tumors (i.e., RB). **(E–F)** Rank–rank hypergeometric overlap (RRHO) analysis comparing differential expression signatures between primary and fetal cells and stem cell-induced RB and fetal cells.

Furthermore, we identified the cell types of 234,689 cells from the retina. By employing the machine learning-based CellTypist tool (model: Human_Developmental_Retina)[47] and known canonical markers of the retina, such as RHO in photoreceptors, we confirmed the cell types. In total, thirteen cell types were identified, including amacrine cells (AC), bipolar cells (bipolars), early glial cells, glial cells (Müller glia), horizontal cells (HC), photoreceptors and progenitors (Photo/Progen), retinal ganglion cells (RGCs), transition zone 1 of progenitor cells (T1) and transition zone 2 of progenitor cells (T2) (**Figure 6C**); the abbreviated names are used hereafter. The data in **Supplemental Figure 7** confirmed that the known canonical markers of each cell type were exclusively expressed, supporting the accuracy of the cell type classification of ours. The identified cell types and proportions aligned well with biological knowledge. For example, cells from the normal adult retina mainly consist of photoreceptors[48], whereas fetal and pediatric cells included a substantial fraction of progenitor cells. In addition, tumor cells (regions in the green dashed line in **Figure 6C**) mainly included progenitors, aligning with our understanding that RB mainly originates from the loss of RB1 precursor cells during early development[49]. Moreover, the abundance of genes encoding relevant RB markers, including DEK and E2F3, is shown in **Figure 6D**. Using GENOAR, we recapitulate known biological mechanisms of RBs without prior knowledge.

On the basis of the confidence in the cell type identification, we further evaluated cell populations by condition and study. **Table 2** lists the identified cell types and numbers of cells by tissue condition (healthy control, CP and tumor) and study set (GSE147979 = samples from normal adults without tumors; GSE166173 = fetus control samples and primary tumor samples; GSE174200 = normal adult control samples and stem cell-induced RB organoids). Although the RB cells from the primary tumor and stem cell-derived organoids were clustered together, their cell type compositions differed substantially. RB cells from primary tumors (from GSE166173) primarily included sensory neurons (photoreceptors, photo/progen, and bipolars) and resident immune cells, such as glia, early glia and microglia. However, stem cell-derived RB organoids present mainly retina-specific neurons, including photoreceptors, photoreceptors/progen, and progenitors, with limited numbers of resident immune cells, including microglia. Among the 55,102 cells from RB organoids (**Table 2**; tumor condition in GSE174200), more than 60% of the cells (n=34,382) were progenitors, and 37% were photoreceptors. In contrast, the native tumor samples included more diverse cell types, including yolk sac-derived resident immune cells. Although the lack of resident immune cells is expected in standard retinal organoids because of their distinct neuroectodermal origin, we identified the presence of microglia (marked number in **Table 1**) and confirmed the presence of characteristic genes of microglia, including AIF1 (**Supplemental Figure 8**). Interestingly, RB cells from distinct microenvironments consistently showed ectopic activation of SALL1 in progenitors and photo/progen. In contrast, no such ectopic activation was detected in progenitors from normal fetal (early developmental stage) or adult retinal cells, indicating that RB organoids partially recapitulate the transcriptional profiles of primary RB. Although the ectopic expression of SALL1 in progenitors and photo/progen requires further experimental validation, GENOA’s tissue aligned multi-conditional cell *atlases* have raised new questions that could lead to novel biological findings.

**Table 2.** Fractions of cell types in the retina by condition and study.

| Cell type | Tissue condition | Cell count | By study set<sup>1</sup> |  |  |
| --- | --- | --- | --- | --- | --- |
|  |  |  | GSE147979<sup>2</sup> | GSE166173<sup>3</sup> | GSE174200<sup>4</sup> |
| Photoreceptors<br>(total number of<br>cells = 78306) | Healthy Control | 35627 | 35627 | 0 | 0 |
|  | CP | 26303 | 0 | 7775 | 18528 |
|  | Tumor | 16376 | 0 | **<u>2342</u>** | **<u>14034</u>** |
| Photo/Progen<br>(total number of<br>cells = 49409) | Healthy Control | 1680 | 1680 | 0 | 0 |
|  | CP | 41079 | 0 | 40808 | 271 |
|  | Tumor | 6650 | 0 | **<u>217</u>** | **<u>6433</u>** |
| Progenitors<br>(total number of<br>cells = 46683) | Healthy Control | 33 | 33 | 0 | 0 |
|  | CP | 7054 | 0 | 7054 | 0 |
|  | Tumor | 39596 | 0 | **<u>5214</u>** | **<u>34382</u>** |
| Bipolars<br>(total number of<br>cells = 21104) | Healthy Control | 10726 | 10726 | 0 | 0 |
|  | CP | 10350 | 0 | 5748 | 4602 |
|  | Tumor | 28 | 0 | 14 | 14 |
| AC<br>(total number of<br>cells = 19423) | Healthy Control | 2470 | 2470 | 0 | 0 |
|  | CP | 16905 | 0 | 16009 | 896 |
|  | Tumor | 48 | 0 | 18 | 30 |
| HC<br>(total number of<br>cells = 5039) | Healthy Control | 489 | 489 | 0 | 0 |
|  | CP | 4543 | 0 | 4302 | 241 |
|  | Tumor | 7 | 0 | 5 | 2 |
| Early Glia<br>(total number of<br>cells = 3218) | Healthy Control | 9 | 9 | 0 | 0 |
|  | CP | 1483 | 0 | 1481 | 2 |
|  | Tumor | 1726 | 0 | 1710 | **16** |
| Microglia<br>(total number of<br>cells = 3067) | Healthy Control | 841 | 841 | 0 | 0 |
|  | CP | 836 | 0 | 555 | 281 |
|  | Tumor | 1390 | 0 | 1220 | **170** |
| Glia<br>(total number of<br>cells = 3031) | Healthy Control | 1088 | 1088 | 0 | 0 |
|  | CP | 1176 | 0 | 174 | 1002 |
|  | Tumor | 767 | 0 | 757 | **10** |
| RGC<br>(total number of<br>cells = 2292) | Healthy Control | 181 | 181 | 0 | 0 |
|  | CP | 1970 | 0 | 1955 | 15 |
|  | Tumor | 141 | 0 | 130 | 11 |
| T2<br>(total number of<br>cells = 1544) | Healthy Control | 2 | 2 | 0 | 0 |
|  | CP | 1542 | 0 | 1542 | 0 |
|  | Tumor | 0 | 0 | 0 | 0 |
| T1/RGC<br>(total number of<br>cells = 799) | Healthy Control | 16 | 16 | 0 | 0 |
|  | CP | 781 | 0 | 781 | 0 |
|  | Tumor | 2 | 0 | 0 | 2 |
| T1<br>(total number of | Healthy Control | 2 | 2 | 0 | 0 |
|  | CP | 772 | 0 | 772 | 0 |
| cells = 774) | Tumor | 0 | 0 | 0 | 0 |
|  | Total | 234689 | 53164 | 100583 | 80942 |

<sup>1</sup>The identifiers of the GEO series (GSE); one GSE presents a unit of research project. <sup>2</sup>Samples from adult deceased donors, <sup>3</sup>Retinoblastoma samples from pediatric patients and matched pairs from fetal samples as CPs. <sup>4</sup>Stem cell-derived retinoblastoma samples for tumors and adult controls as CPs.

Given that the ectopic activation of genes was independently observed in both primary and organoid tumors, we next considered whether this reflected a broader similarity between the two models (native tumors and RB organoids). Thus, we compared their transcriptional signatures using the rank–rank hypergeometric overlap (RRHO) approach[50]. The RRHO approach enables a threshold-free comparison of changes in gene expression using the listed p values and effect sizes (i.e., fold changes (FC)) of genes. Here, we computed the FCs of genes using pseudobulk transcriptomic profiles across all cells and progenitor cells. To calculate FCs, we counted pseudobulk gene expression from all cells and progenitors in primary RBs (tumor condition in GSE166173), fetal control cells (CP condition in GSE166173), and organoid RBs (tumor condition in GSE174200). Afterward, we calculated the rank of the genes according to the sign(log2FC) × −log10 P value from the primary RB vs fetal control comparison and organoid RB vs fetal control comparison. Then, the R package of RRHO2[51] was used to visualize the degree of concordance among the ranks of the genes using a quadrant plot for all cells and progenitors (**Figure 6E, F**). In the RRHO quadrant plots, the X-axis indicates the rank of the genes from the organoid RB vs. fetus control comparison. The y-axis presents the rank of the genes from the primary RB vs. fetus control comparison. The color denotes the statistical confidence based on the significance of the overlap (−log10 adjusted P value). In summary, the red color in the upper-right/lower-left regions in the quadrant plots indicates concordance (up-up/down) between primary and organoid tumors, whereas results in the upper-left/lower-right regions are discordant. The RRHO results for all the cells presented a limitation of the RB organoids because of a lack of shared concordances (blue in the first quadrant for the upper-right panel, **Figure 6E**). However, the results from progenitors were strongly concordant (**Figure 6F**). In addition, we evaluated the concordance of cell cell interactions and the microenvironment by highlighting known ligand receptor genes (white circles) using ligand receptor genes from CellPhoneDB[52]. Interestingly, the expression of a ligand, IL23A, was strongly correlated between progenitor cells in RB organoids and primary RB (**Figure 6F**, first panel, red colored region; -log(P)≈300), which may indicate that the inflammatory microenvironment is shared between native and organoid tumors. However, structure-associated ligands of progenitor cells, such as COL2A1, showed only marginal concordance (**Figure 6F**, second panel, blue colored region; -log(P)≈0.0) because of the limited architecture in the organoid models. These distinct patterns indicate that RB organoids recapitulate the inflammatory signals in the microenvironment; however, they have intrinsic limitations in terms of mimicking the spatial tissue niche (**Supplemental Data 5** list the genes shown in the quadrant plot).

Taken together, these findings demonstrate that GENOAR can systematically organize diverse transcriptional profiles from large-scale scRNA-seq data and subsequently facilitate data-driven hypothesis generation.

## Discussion

In this study, we established a general framework in which single-cell transcriptomics *atlases* were constructed in a scalable and AI-compatible manner. Our developed system, named GENOAR (the global engine for navigating the Omnicell space via agent-based research), covered approximately 54 million cells for diverse downstream tasks using AI agents and parallel computing. The three key components of GENOAR highlight the originality of our study. The first component is the reproducible organization of single-cell transcriptomic metadata in the GEO by aligning AI agents with a concept-based corpus using the UMLS under the guidance of human experts. The second component is the demonstration of the on-demand construction of single-cell atlases in *Mus musculus* without consortium-like updates or the manual integration of scRNA-seq data. The third component involves the application of parallel computing technology, enabling the petabyte-scale preprocessing of scRNA-seq data and allowing diverse downstream tasks for human researchers and AI agents. In addition, the performance of the AI agents in GENOAR was validated on the basis of the RAGAS score. Thus, GENOAR is a reusable framework, and the automated construction of analysis-ready datasets mitigates technical challenges in the AI era. Because AI models inherently use substantial data as the use of millions of parameters, GENOAR’s continuous supply of AI-compatible scRNA-seq data enables iterative, ongoing model updates as the amount of raw data increases.

The unprecedented success of LLMs and generative models has revolutionized single-cell biology, enabling the development of foundation models(7) for diverse downstream tasks. However, these models face key limitations. Most are not designed for zero-shot prediction; they require fine-tuning with an appropriate dataset for each task, and they are restricted mainly to human data. Although zero-shot modeling in single-cell biology has recently advanced to include UCEs [54], effective, broad-scale training on nonredundant scRNA-seq data remains an unsolved challenge. To address these constraints [53], a scalable approach to single-cell atlas construction, coupled with organized metadata, is essential. Ideally, the on-demand construction of single-cell atlases that are updated as new raw data become available, generalize across multicellular organisms, and pair standardized metadata with analysis-ready datasets should be constructed. GENOAR enables the systemic construction of single-cell atlases for different species; then, by packaging the complex pipeline into a Docker image, users can execute these multistep processes without the need for specialized hardware or infrastructure expertise. In addition, with the AI librarian and AI disease nomenclaturist in GENOAR, users can seamlessly prioritize scRNA-seq data to fine-tune given foundation models. Thus, GENOAR makes systematic, integrative analyses of large-scale transcriptional data more broadly accessible allowing democratization of single-cell atlases analysis. As initiatives such as CZ CELLxGENE and the Human Cell Atlas[5] have increased access to harmonized human datasets, GENOAR complements this open-science movement by extending this access to diverse multicellular species via an AI-assisted open-source framework.

In addition to the contributions of our study, several limitations should be addressed. In the present study, we tailored our preprocessing pipeline for scRNA-seq data based on a fixed modality (10X Genomics Inc., Chromium) with specific read-depth criteria. Although the presented framework targeted the most popular sequencing technology in the single-cell field, the data-processing pipeline could be further expanded. Additionally, owing to the medical-focused nature of the utilized UMLS, the computational efficiency of AI disease nomenclaturist must be improved to evaluate diverse organisms in the context of nonhuman diseases and organism-specific tissues. We also acknowledge that the current version of GENOAR could be developed further. First, the source metadata of the GEO could be further organized, which remains loosely controlled and requires orchestration by human experts. For example, in the current version, several cancer studies were identified via the ‘tissue’ field based on the information submitted to the GEO by the researcher. Although we developed a generalized organization approach for studies with unstructured metadata, the metadata analysis could be further improved. However, to our best knowledge, GENOAR initially enables the construction of single-cell *atlases* in an on-demand manner from scratch, whereas previous works focused on either metadata organization[55] or data integration[10]. Moreover, by releasing the source code of GENOAR, this limitation should be addressed in a timely manner through the community of users (https://github.com/cxinsys/genoar).

From the development of GENOAR, several insights can be deduced. First, human experts and AI agents each have advantages. Although we implemented AI agents and a RAG-based AI-compatible dataset, the guidance of human experts and structured domain-specific knowledge, including that from the UMLS, improved the performance of the AI agents in GENOAR. Second, the computing environment itself is beginning to shape the bioinformatics field, including AI, in a distinct way; both GPU performance and lower-level technology need to be considered. The computational challenges in the construction of the single-cell atlas differed from the challenges observed in general-purpose HPC settings, which mainly involves optimizing the raw computational throughput (e.g., FLOPS). Although file I/O (i.e., the sheer volume of single-cell files) was the primary constraint during atlas construction, downstream analyses, including data integration in our RB analysis, were limited mainly by GPU memory. Thus, we anticipate that the next technical challenge to be addressed in single-cell biology is memory optimization via multiple GPUs or hybrid computations with CPUs and GPUs. Finally, industry-driven AI applications in biomedical science reflect both hype and hope[56, 57]. The remarkable scale and downstream performance of existing methods increase our hope in this technology. However, the limited transparency in model-building code and the lack of rigorous agent evaluations make it unclear whether performance improvements reflect genuine agentic accuracy or simply data scale. Thus, this highlights the potential of AI-driven biological data resources and the expectation of reproducible results for each researcher. Accordingly, GENOAR is released as fully open-source software, including not only its output datasets but also its data-building modules and the quantitative validation of its retrieval components.

In addition to atlas construction and the rigorous performance evaluation, GENOAR can be applied in downstream analysis tasks, as demonstrated by our RB case study. The objectives of our RB analysis were to examine the quality of the data integrated by GENOAR and the feasibility of executing downstream tasks, including hypothesis generation. As previously described, RB is a pediatric cancer that is peculiarly driven by RB1 deletion in retinal progenitors. Thus, without GENOAR, such integrative, cross-model (i.e., primary vs. organoid RB) comparisons would have demanded considerable time and computational expertise. Consistent with the cartographic definition of *atlases*, GENOAR provided an integrated RB atlas spanning diverse conditions, including adult healthy retinas. This integration emerged naturally from the label-based and AI agent-assisted data aggregation process rather than deliberate manual curation of data. Consequently, this integrative capacity allowed us to resolve SALL1 expression, which is specific to fetal tumor cells and is absent in both the normal adult retina and the fetal retina (Supplemental Figure 8). Taken together, these findings suggest that the generalized presentation of cell atlases across conditions may facilitate the development of biological hypotheses.

Nobel laureate Sydney Brenner noted the necessity of developing virtual cells, as they represent the fundamental units of life, making them a predictive model for the next era of biology. He also indicated the need for such models to be computationally efficient. We believe that GENOAR paves the way toward this vision. We presented an advanced approach for harmonizing the annotation and industry-level preparation of single-cell transcriptomic data in an AI-compatible manner, enabling the scalable building of models for cell biology.

## Materials and Methods

### Retrieval of submitted metadata from the GEO and analyses

The first component of GENOAR is the retrieval of the metadata submitted to the GEO. From the web page of the GEO (https://www.ncbi.nlm.nih.gov/geo/), GENOAR programmatically collected a list of GSEs (the identifier of one research project in the GEO) on the basis of user-defined keywords (details in **Supplemental Data 6**).

### Construction and validation of AI agents in GENOAR

To construct a RAG-based agent (i.e., an AI librarian), we collected scRNA-seq metadata from the NCBI GEO, including the SRR identifier of the Sequence Read Archive (SRA) and samples from human and mouse studies. The raw metadata included diverse biological annotations, such as tissue types, disease states, cell types, and experimental conditions. Core metadata fields— including Run ID, Series ID, BioSample ID, Sample Name, and standardized biological descriptors—were prepared using the UMLS. In this step, we also employed an additional AI agent (disease nomenclaturist) to guide the disease-to-disease relationship. We implemented a MCP-based autonomous agent powered by Google Gemini for the iterative integration of knowledge from the UMLS. Human experts (research experiences 3∼15 years) guided the disease and diagnosis identification and assigned a disease classification level. All the integrated disease names given by the AI disease nomenclaturist were manually validated. Further details of database indexing and storage systems are described in Supplemental Data 6.

### Evaluation of the performance of the AI librarian (RAG-based agent)

We evaluated the performance of the RAG retrieval system (AI librarian) using a comprehensive framework that combines RAGAS metrics and traditional information retrieval (IR) metrics to assess both semantic relevance and ranking quality. The RAGAS scores were computed as described in Supplemental Data6.

### Parallelized preprocessing of scRNA-seq data and exploration in GENOAR

After we retrieved and normalized the metadata from the GEO, we fetched the raw scRNA-seq data and then converted these data into FASTQ format. Using our developed dynamic workload pipeline, all the raw data were processed into an analysis-ready format (HDF5/h5) for downstream analysis, including data integration and scaling. We released a web interface and Ducker containers for the convenient exploration of the established single-cell atlases (https://github.com/cxinsys/genoar.git) (Supplemental Data 6).

### Identification of multicondition single-cell atlases

To retrieve samples representing multiple conditions for the matched tissue, we applied a keyword-based approach to the standardized metadata of GENOAR. Details of the identification of samples are presented in Supplemental Data 6.

### GPU-aided analysis of scalable atlases of CPs, tumors and healthy controls

Count matrices from the classified CP, tumor, and healthy control samples were merged and processed using Scanpy and RAPIDS-SingleCell[58], a GPU-accelerated single-cell analysis library. A combined tissue-condition label was constructed by integrating tissue type and group identity for downstream visualization.

Quality control was performed by filtering cells on the basis of the detected gene counts (<5000∼7000) and mitochondrial gene fractions (<20%), followed by the removal of genes whose expression was detected at low levels in our previous work[59]. All GPU-accelerated steps were executed on an NVIDIA GPU with RMM-managed memory optimization. The overall pipeline for GPU-accelerated single-cell transcriptomics was presented in the web (https://github.com/scverse/rapids-singlecell). In the CPU run, a single node with 64 AMD EPYC virtual cores and 512 GB of RAM was used; in the GPU run, one NVIDIA H100 with 80 GB HBM3 (host CPU: AMD EPYC 9654) was used (details in **Supplemental Data 6**).

### Declaration of generative AI and AI-assisted technologies in the writing process

During the preparation of this work, the authors used AI in very limited contexts to improve writing efficiency, such as to identify typos and address grammar issues. After using this tool/service, the authors reviewed and edited the content as needed and take full responsibility for the content of the publication.

## Resource availability

### Lead contact

Further information and requests for resources should be directed to and will be fulfilled by the lead contact, Hyojung Paik and Daewon Lee.

### Material availability

No new materials were generated in this study.

### Code availability

The original code is also available at https://github.com/cxinsys/genoar.git and https://github.com/cxinsys/genoar.

### Data and web resources

All the processed datasets are available for humans and AI models at https://genoar.ai.

## Acknowledgment

This work was supported by the Korea Institute of Science and Technology Information (KISTI) (K26L1M2C3). This work was supported by National Research Foundation of Korea (NRF) grants funded by the Korea government (MSIT) (RS-2025-16065483; RS-2025-02315385). This research was also supported by the project of Building and Operating the Korea-Bio Data Station (K-BDS) Platform Building and Tool Development (NRF-2021M3H9A2030520). This work was supported by the National Supercomputing Center with supercomputing resources, including technical support. This research was also supported by the Basic Science Research Program through the National Research Foundation of Korea (NRF), which is funded by the Ministry of Education (NRF-2017R1A6A1A03015713).

## TABLE OF CONTENTS FOR SUPPLEMENTARY MATERIALS

**Supplemental Figure 1.** On-demand construction of a large-scale single-cell transcriptomics atlas of mice via GENOAR.

**Supplemental Figure 2.** Matched results between manual disease/tissue naming and the GENOAR classification.

**Supplemental Figure 3.** Classification results of single-cell transcriptomic data using the AI disease nomenclaturist in GENOAR.

**Supplemental Figure 4.** Detailed results of the on-demand classification of the disease type on the basis of mouse data.

**Supplemental Figure 5.** Identification of tumors, healthy controls and matched control pairs (CPs) in GENOAR.

**Supplemental Figure 6.** Batch correction and detailed t-SNE clustering results of retinal cells by study and sample.

**Supplemental Figure 7.** Confirmation of the cell type predictions using known marker genes of retinal cells.

**Supplemental Figure 8.** Detailed t-SNE clustering results and microglia-related gene expression in retinal cells.

**Supplemental Data 1.** A manually integrated metadata set of single-cell transcriptomic data from the GEO (Jan. 1, 2010 to Dec. 31, 2022).

**Supplemental Data 2**. Expert validation set for the AI agent.

**Supplemental Data 3.** CPU and GPU execution times (seconds) for the single-cell processing pipeline.

**Supplemental Data 4.** List of samples for the analysis of the multicondition tissue atlases.

**Supplemental Data 5.** List of concordant genes in the RB analysis.

**Supplemental Data 6.** Details of methods.

## Author contributions

Hyojung Paik: Conceived the main idea, acquired the data, wrote the manuscript draft, designed the bioinformatic analysis, developed the agents, supervised the overall project, prepared all the figures, and acquired funding.

Tae Lyun Ko: Carried out the bioinformatics analysis, developed the agents, developed the source code, acquired the data, prepared the figures, and developed the website.

Dogyun Kim: Carried out website development, prepared the graphical abstract, and performed the Dockerization of the source code

Hyeonjae Lee: Supported the Dockerization of the source code

Taeyoung Lee: Supported the bioinformatics analysis and prepared the main figures. Dongmin Shin: Performed the Dockerization of the source code

Marina Sirota: Supported the source code validation Boris Oskotsky: Supported the source code validation Tomiko Oskotsky: Supported the source code validation

Daewon Lee: Overall supervision of the Dockerization of the source code and web development

