## Supplementary figures and images for "GENOAR: Global Engine for Navigating the Omnicell space via Agent-based Research"

### SFig1.jpg

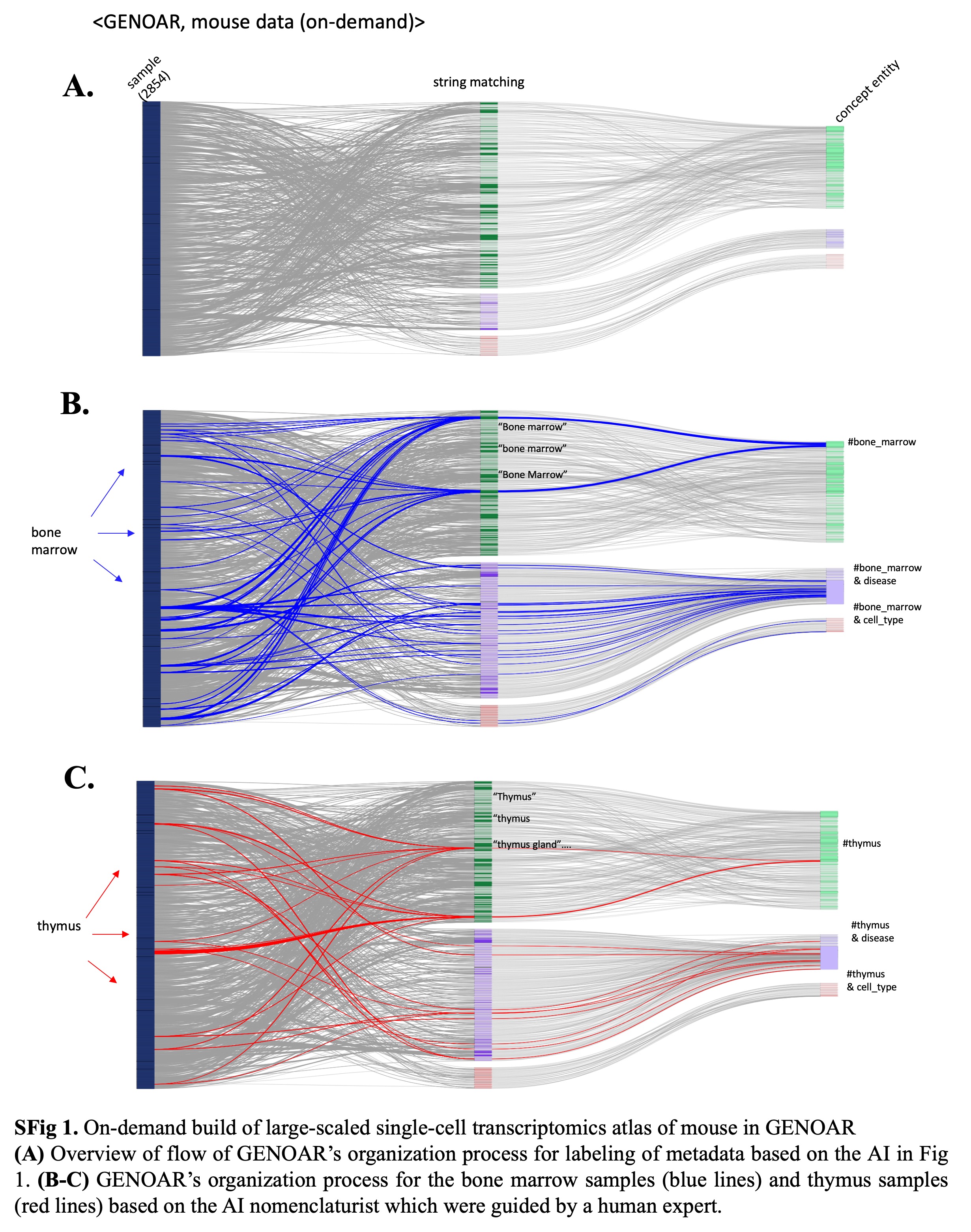

### SFig2.jpg

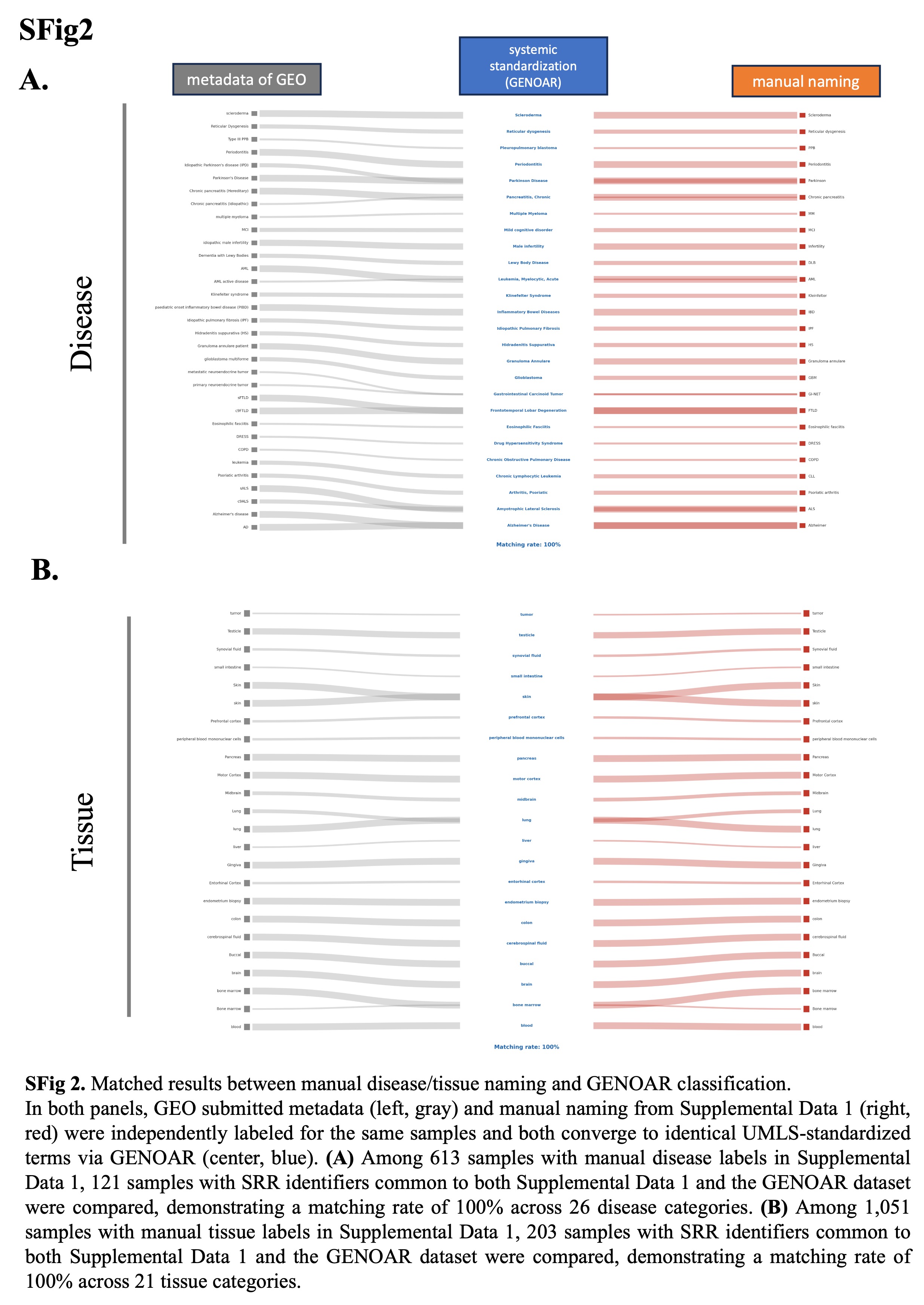

### SFig3.jpg

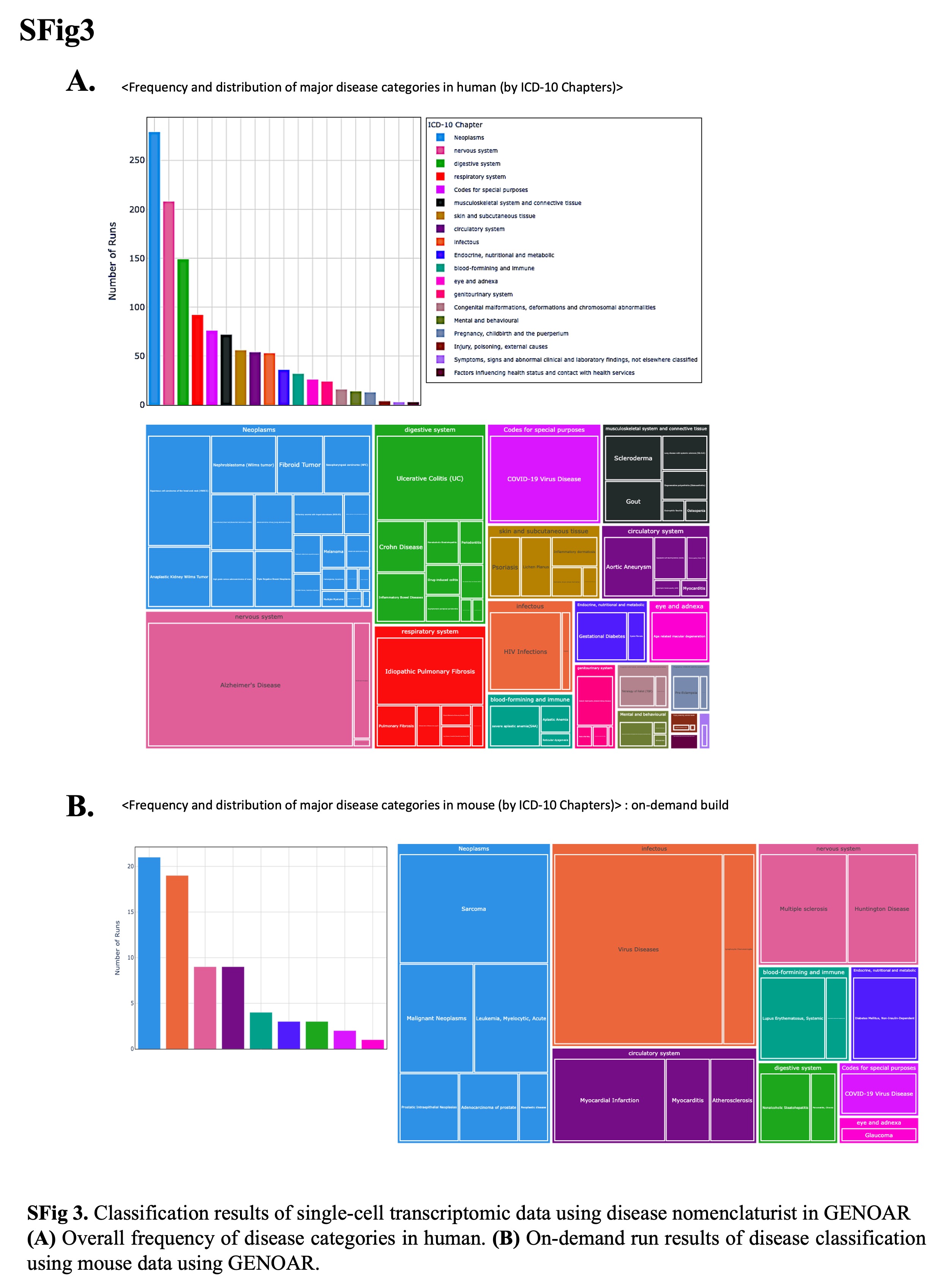

### SFig4.jpg

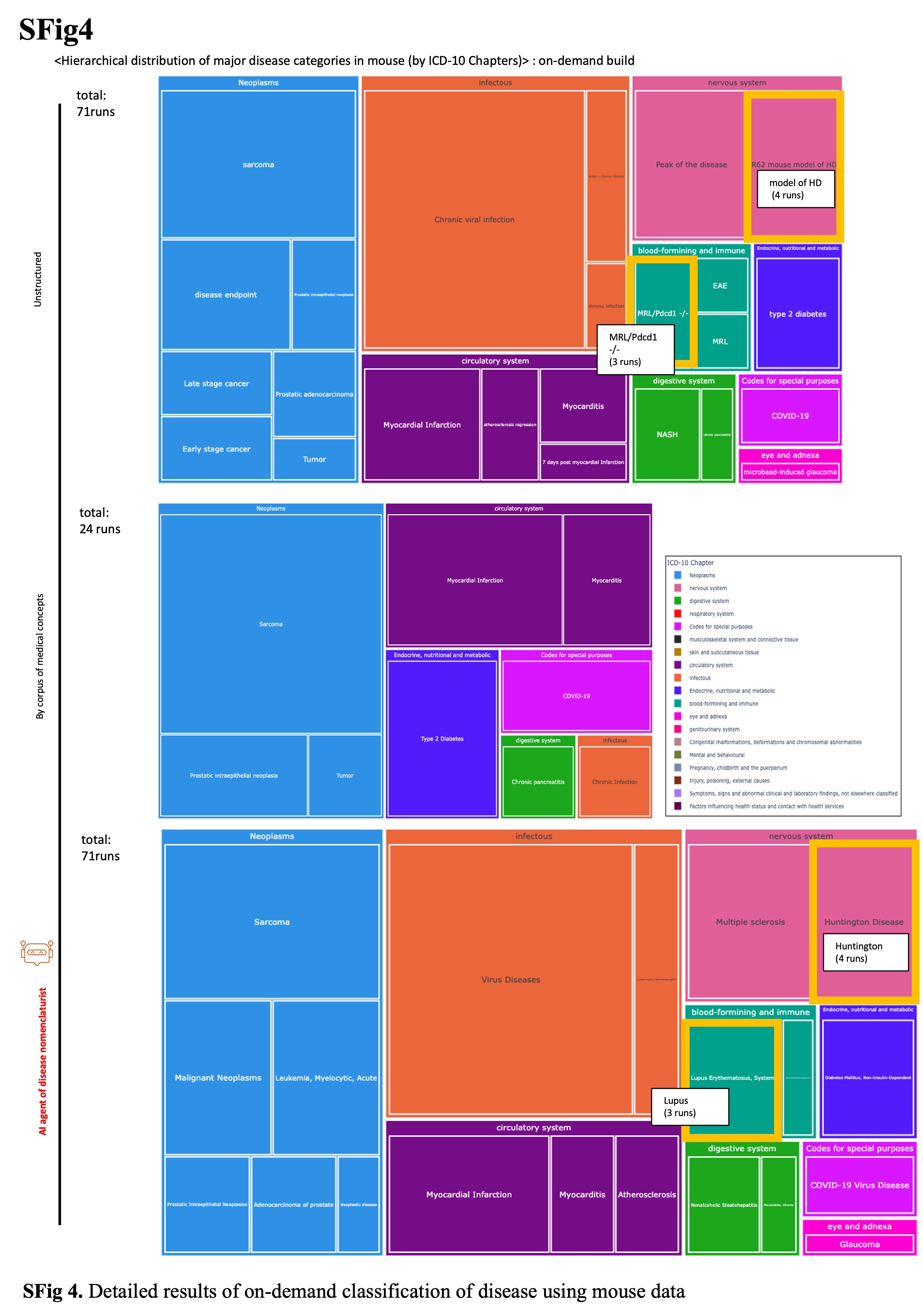

### SFig5.jpg

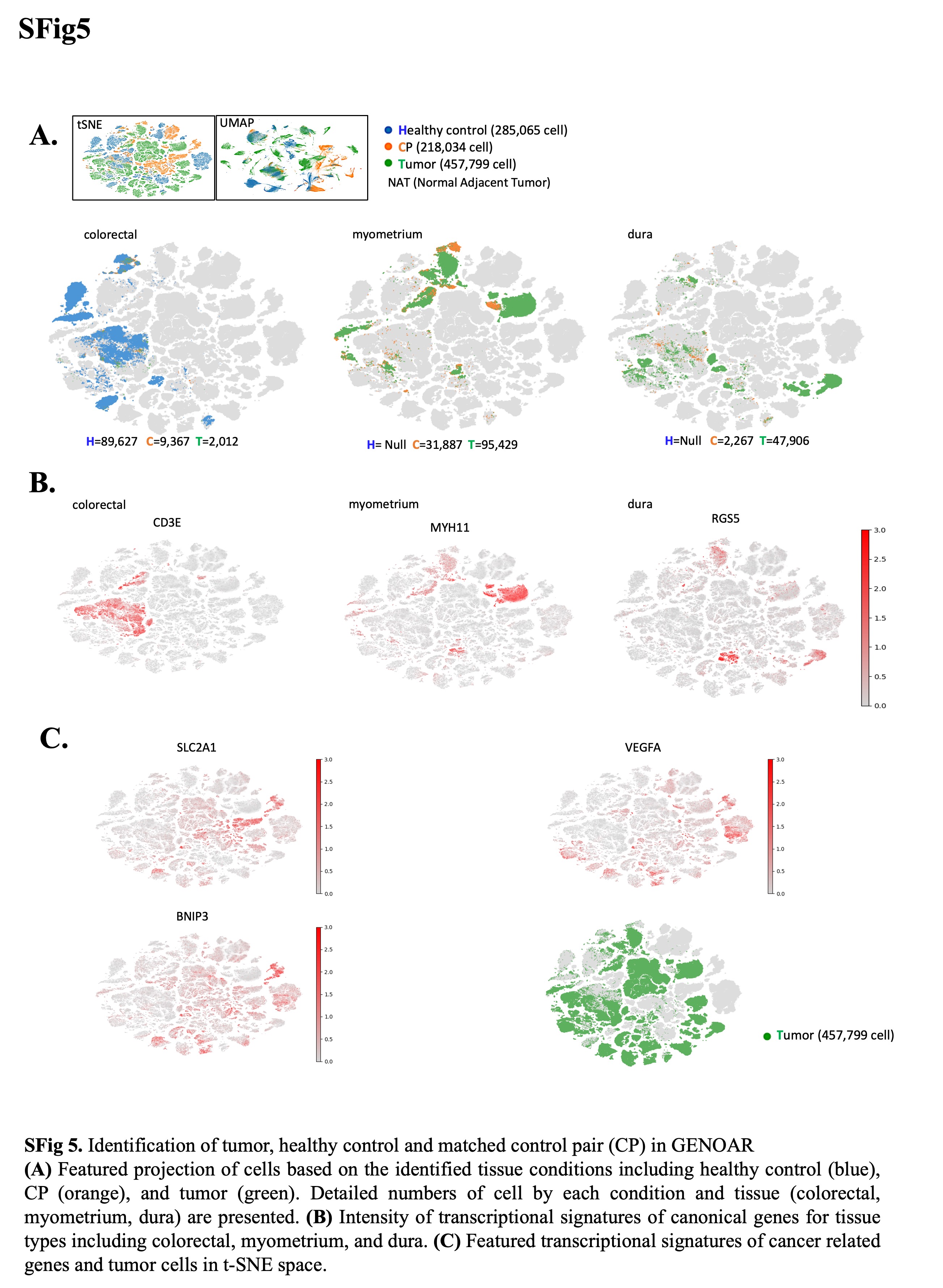

### SFig6.jpg

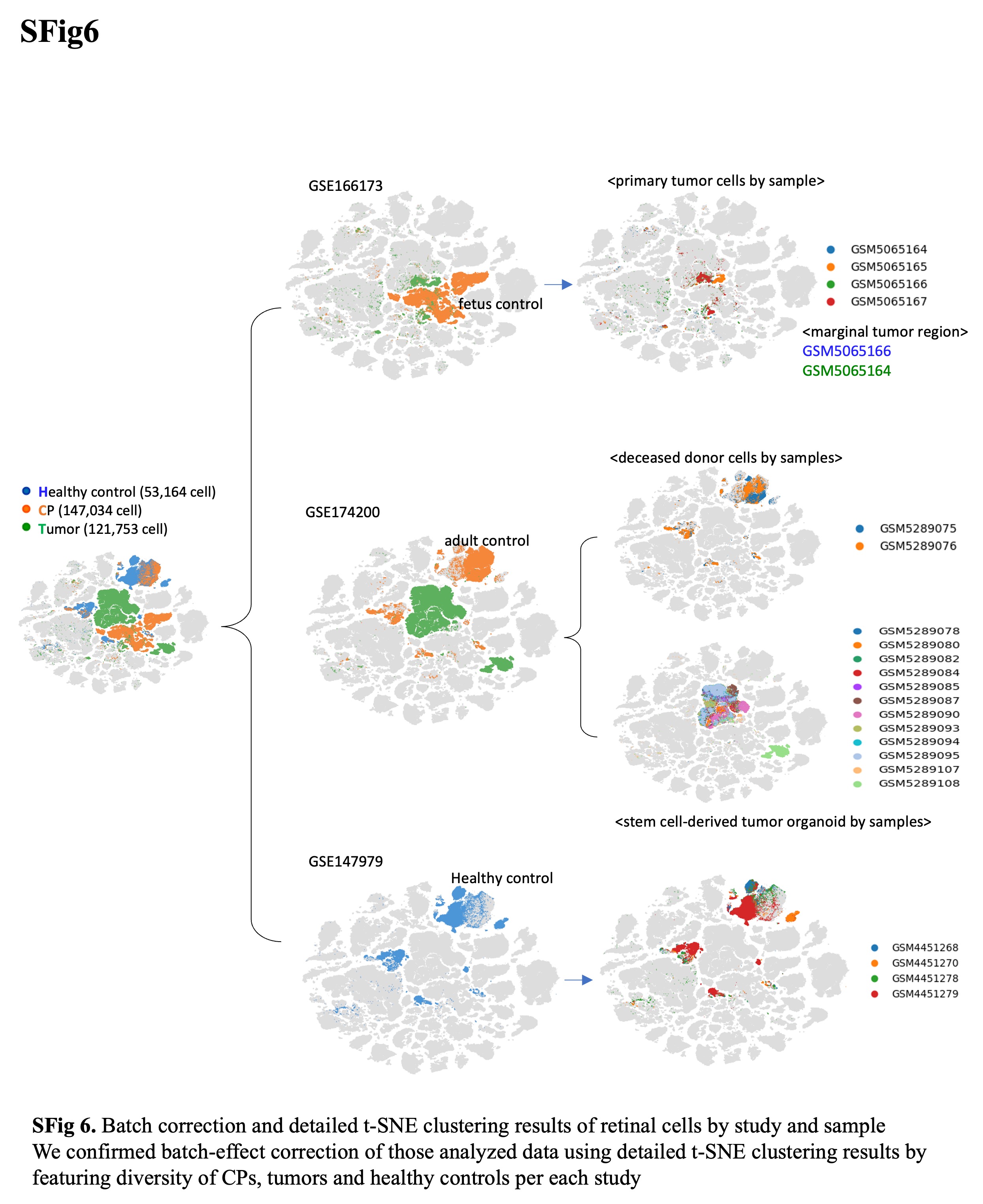

### SFig7.jpg

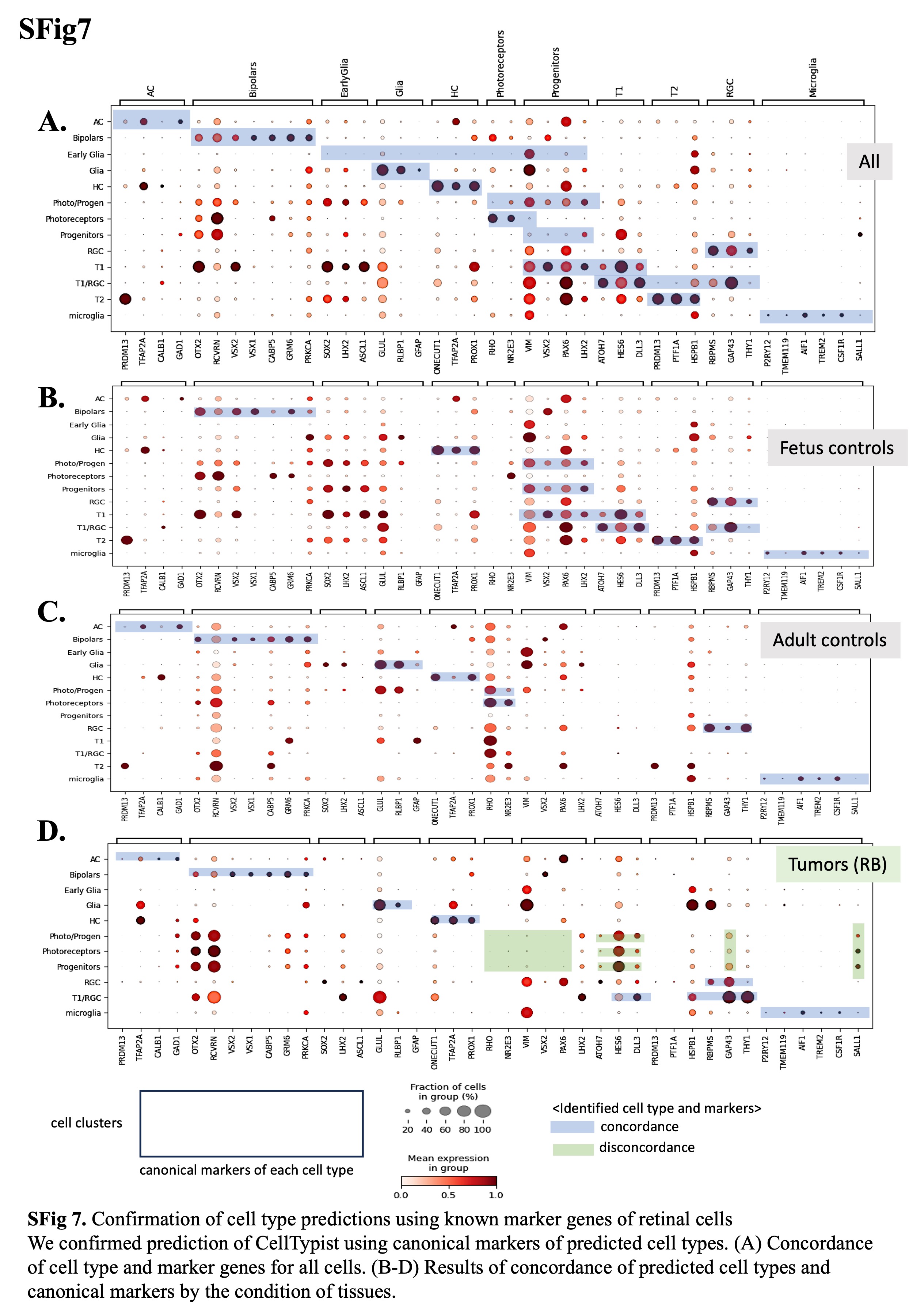

### SFig8.jpg

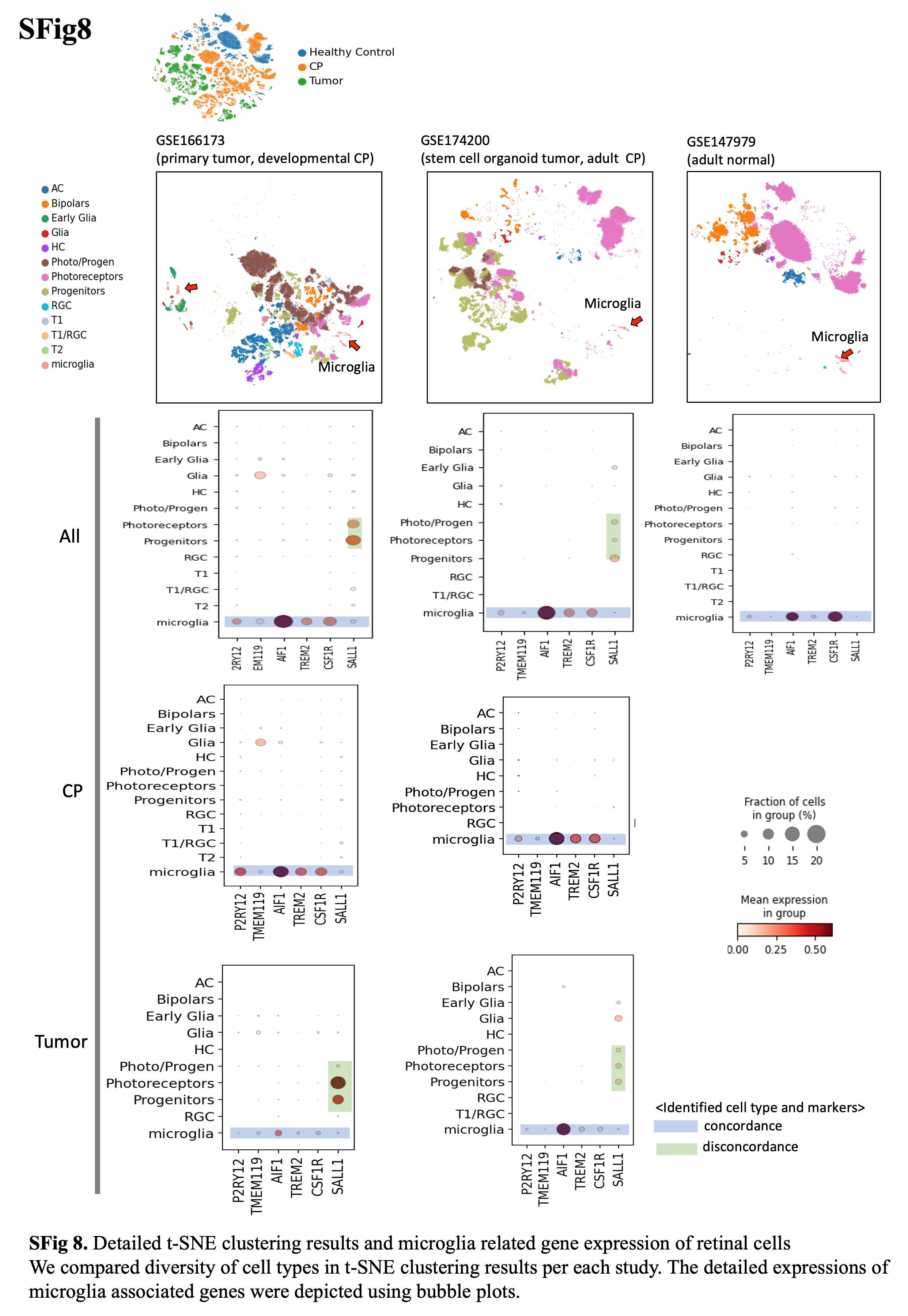
